# PI3Kδ promotes T cell effector differentiation and plasticity during chronic infection

**DOI:** 10.64898/2025.12.19.695585

**Authors:** Andrea C. Pichler, Jennifer L. Cannons, Dominic P. Golec, Julie M. Reilley, Dan Corral, Eduard Ansaldo, Qin Xu, Subrata Paul, Paul Schaughency, Francisco A. Otaizo-Carrasquero, Stacie M. Anderson, Anshu Deewan, Dorian B. McGavern, Pamela L. Schwartzberg

**Affiliations:** Laboratory of Immune System Biology, National Institute of Allergy and Infectious Diseases, Bethesda Maryland, USA; Laboratory of Host Immunity and Microbiome, National Institute of Allergy and Infectious Diseases, Bethesda Maryland, USA; Integrated Data Sciences Section, Research Technologies Branch, National Institute of Allergy and Infectious Diseases, National Institutes of Health, Bethesda, MD 20892, USA; National Human Genome Research Institute, Bethesda Maryland, USA; Viral Immunology and Intravital Imaging Section, National Institute of Neurological Diseases and Stroke, Bethesda Maryland, USA

## Abstract

Persistent antigen exposure in chronic infections and cancer leads to a progressive state of T cell dysfunction known as exhaustion, which represents a major barrier to effective immune control, but allows antigen-specific T cells to persist. Understanding signaling pathways that mitigate exhaustion and reinvigorate CD8^+^ T cell effector function is a key goal for immunotherapeutic strategies. Here, we show that an activating mutant of phosphoinositide-3-kinase δ (PI3Kδ) led to a reduction of FoxO1-dependent TCF-1^+^ stem-like progenitor CD8^+^ T cells that are required for sustaining antigen-specific T cells in response to chronic viral infection. Nonetheless, mice expressing activated PI3Kδ maintained CD8^+^ T cell responses that were skewed instead towards effector-like cells in a FoxO1-independent manner, associated with an amplified IL-21-STAT3 response axis and improved viral control. Activated PI3Kδ limited TOX expression, prevented epigenetic changes associated with T cell exhaustion, and promoted effector differentiation and function from both progenitor stem-like cells and cells with an exhausted phenotype. Together, this work uncovers a key role for PI3Kδ activation in shaping the balance and plasticity between effector function and exhaustion while promoting T cell persistence during chronic infection, providing insight for immunotherapeutic strategies.

## Introduction

CD8^+^ T cells play a key role in providing protection against infection and cancer. However, during chronic infection and cancer, prolonged antigen exposure drives a state of progressive T cell hyporesponsiveness, known as “T cell exhaustion”. Exhausted T cells (Tex) are characterized by the expression of inhibitory receptors such as PD-1, Tim3 and LAG3 that dampen T cell signaling and effector function ^1, 2^. During exhaustion, antigen-specific T cells are maintained by a small subset of stem-like progenitor (Tex^PROG^) cells, that is defined by and requires the expression of the transcription factor TCF-1 and which transcriptionally resemble memory T cells in acute infection. Tex^PROG^ cells both self-renew and give rise to the more dysfunctional TCF-1^-^ terminally Tex cell population ^3, 4, 5, 6, 7^. Exhausted T cells also upregulate TOX, a key epigenetic and transcriptional regulator that is required to establish the T cell exhaustion program and enable Tex cells to persist ^8, 9, 10, 11, 12^. TOX-dependent changes in chromatin accessibility are maintained in Tex cells even after antigen withdrawal, a phenomenon known as “epigenetic scarring”, which prevents recovery of function of terminally exhausted (Tex^TERM^) cells due to a lack of epigenetic plasticity ^13, 14^. Although the loss of responsiveness of CD8^+^ T cells during chronic infection ultimately leads to decreased immune control, it allows antigen-specific T cells to persist while protecting the host from immunopathology ^1, 2^. Because T cell exhaustion hinders resolution of infection or tumor control, understanding the signaling events that govern the differentiation and maintenance of T cells under conditions of exhaustion is critical for designing more effective immunotherapies.

Mapping the dynamics of T cell signaling has revealed how specific pathways (e.g. PD-1, NFAT, m-TOR) modulate exhaustion and how their manipulation can reinvigorate T cell functions ^1, 15, 16, 17^. PI3Ks are lipid kinases that integrate signals from antigen, co-stimulatory, cytokine, growth factor and other receptors and play important roles in T cell activation, survival and differentiation; PI3Kδ is the major isoform expressed by T lymphocytes ^18^. Activated PI3Kδ syndrome type 1 (APDS1), caused by heterozygous activating mutations affecting PI3Kδ, is associated with immune-dysregulation with recurrent respiratory infections, enteropathy and an inability to clear certain chronic infections ^19, 20^. Intriguingly, many molecules that affect T cell exhaustion, including FoxO1, Bach2, KLF2 and cytokines are regulated by PI3K ^21, 22, 23, 24, 25, 26^. Recent data indicate the importance of PI3K signaling in effector and memory differentiation during acute infection, with activated PI3Kδ impairing expression of TCF-1 and central memory formation ^27^. Moreover, inhibition of PI3Kδ in culture can increase TCF-1^+^ cells, which are critical for sustained antitumor and antiviral responses during exhaustion ^28, 29, 30, 31^ and *in vivo*, can decrease T_REG_ function in cancer ^32, 33^. Nonetheless, PI3K is also important for CD8^+^ effector T cell function ^27, 33, 34, 35^; how PI3Kδ activation in T lymphocytes affects their fate during chronic infections remains largely unexplored.

To address these outstanding issues, we used a mouse model of APDS1, *Pik3cd*^E1020K/+^ mice, that express a heterozygous activating mutation of the endogenous *Pik3cd* gene^36^, which we challenged with Lymphocytic choriomeningitis virus clone 13 (LCMV cl13), a prototypical chronic viral infection ^37^. *Pik3cd*^E1020K/+^ mice exhibited higher viral titers and increased mortality in the first week post-infection (p.i.), yet surprisingly, survivors recovered faster with better viral control than WT mice. Although *Pik3cd*^E1020K/+^ mice had fewer TCF-1^+^ Tex^PROG^ cells secondary to FoxO1 inhibition, LCMV antigen-specific cells still persisted. Instead, activated PI3Kδ drove expansion of a CX_3_CR1^+^ effector-like (Tex^EFF-LIKE^) cell population in an IL-21-dependent manner, associated with increased IL-21 producing T follicular helper (T_FH_) cells, elevated responsiveness to IL-21 including increased cell survival, and improved viral control. Notably, CD8^+^ T cells in chronically-infected *Pik3cd*^E1020K/+^ mice had reduced TOX expression, lacked changes associated with epigenetic scarring and showed increased effector function and generation of CX_3_CR1^+^ Tex^EFF-LIKE^ cells upon cell transfer of either Tex^PROG^ or Tex^TERM^, arguing that PI3Kδ activity promotes cellular plasticity and uncouples persistence from progenitor cells during chronic infection. Our results suggest that PI3Kδ plays a dual role during chronic infection, acting as a gatekeeper that promotes effector cell differentiation and function under conditions of exhaustion.

## Results

### Increased mortality yet enhanced viral control during chronic viral infection in mice expressing activated PI3Kδ

To assess the effects of activated PI3Kδ on exhaustion during chronic infection, we infected *Pik3cd*^E1020K/+^ mice or littermate wild-type (WT) controls with LCMV cl13. Unlike WT mice, approximately 50% of *Pik3cd*^E1020K/+^ mice died by day 10 (d10) post infection (p.i.) (**Fig.1A**), with increased clinical scores and elevated serum IFNγ and TNFα, peaking at d8 p.i. compared to WT (**Fig.1B-D**). Nonetheless, surviving *Pik3cd*^E1020K/+^ mice recovered faster than WT littermates, with more rapidly decreasing clinical scores (**Fig.1B**) and increasing weight gain (**Fig.1C**). On d8 p.i serum titers were similar to WT (**Fig.1E**). However, further evaluation revealed that viral titers were significantly increased in some tissues of *Pik3cd*^E1020K/+^ mice early post-infection (d4 p.i.) (**Fig.1F and Extended Data Fig. 1A**), but by two weeks, were lower compared to WT littermates, even though the virus still persisted in all tissues examined, consistent with a chronic infection (**Fig.1F and Extended Data Fig.1A**). These phenotypes were specific to chronic infection in *Pik3cd*^E1020K/+^ mice, as we observed no differences in survival, clinical scores and weight loss in response to acute infection with LCMV Armstrong, compared to WT littermate controls (**Extended Data Fig.1B-D**). Thus, *Pik3cd*^E1020K/+^ mice showed premature lethality and increased early inflammatory immune responses to chronic infection, yet improved viral control at later timepoints.

**Figure 1.**
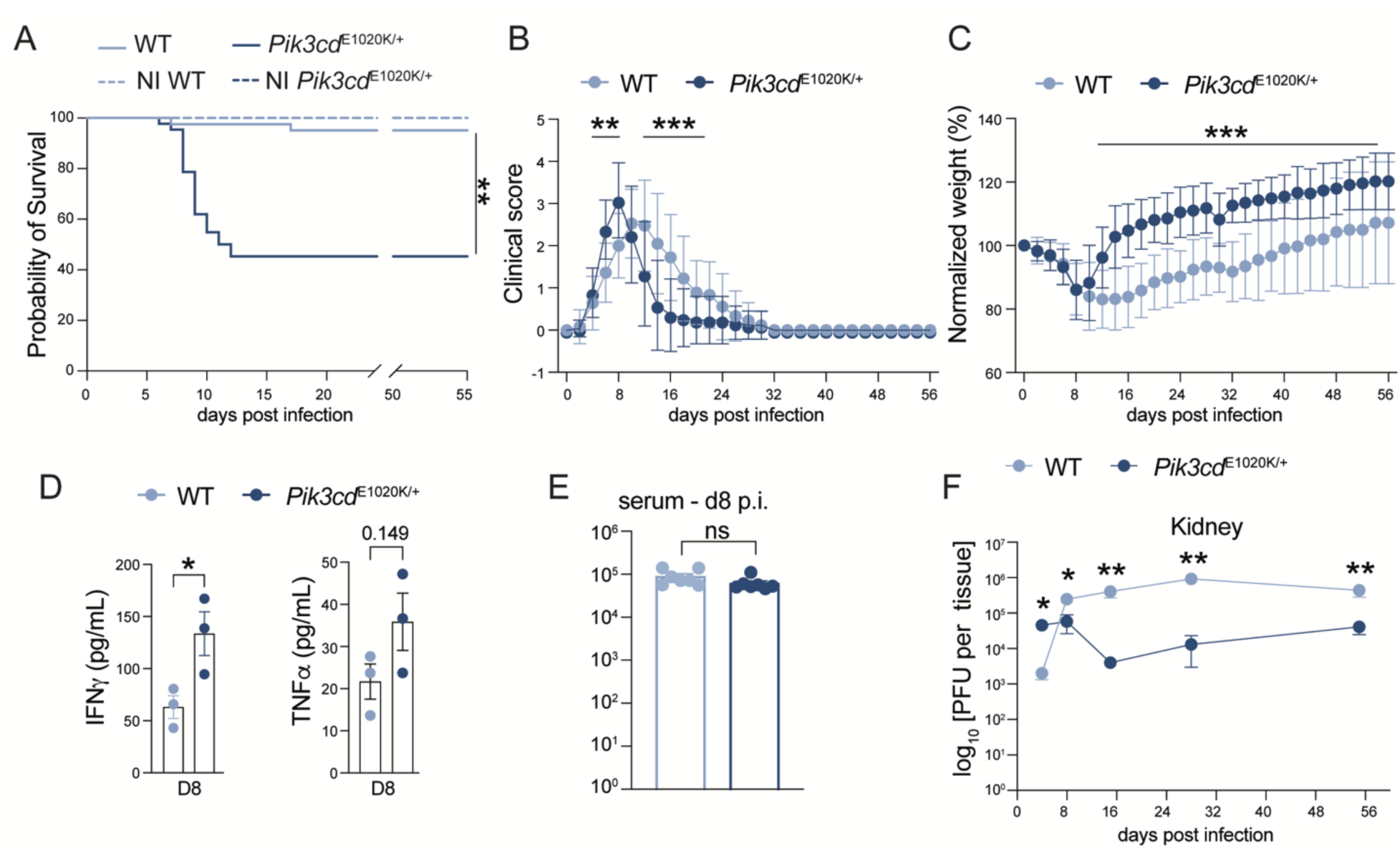
Activated PI3Kδ changes disease outcome during chronic infection. C57BL/6 WT and *Pik3cd*^E1020K/+^ mice were infected i.v. with 2x10^6^ PFU LCMV cl13. Mice were monitored for weight, clinical score and survival**. (A)** Kaplan Meyer survival curve. **(B-C)** Graphs showing clinical score **(B)** and normalized weight **(C)**. **(D)** IFNγ (left) and TNFα (right) in serum d8 p.i. **(E-F)** Plaque forming units (PFU) in serum on d8 **(E)** and kidney **(F)** at indicated time points. In **(B)** and **(C)** each dot represents a pool of 43 (d0-8 WT), 42 (d0-8 *Pik3cd*^E1020K/+^), 20 (d10-30 WT), 18 (d10-30 *Pik3cd*^E1020K/+^), 18 (d32-60 WT) and 17 (d32-60 *Pik3cd*^E1020K/+^). In **(F)** each dot represents pools of 7, 7, 8, and 10 mice on d4, 8, 15, 28 and d55 p.i. in WT mice respectively and 8, 7, 7 and 6 mice on d4, 8, 15, 28 and d55 p.i. in *Pik3cd*^E1020K/+^ mice, respectively. In **(D and E)** each dot represents one mouse, data are representative of two independent experiments (**D),** or are pooled from 2 experiments (**E)**. Statistical differences were determined by Mantel-Cox test **(A)**, one-way ANOVA **(B-C and F)** and Student’s t-test **(D and E)** *p < 0.05, **p < 0.01, ***p < 0.001.

### Activated PI3Kδ drives expansion of effector-like Tex cells at the expense of Tex^PROG^ cells

Because CD8^+^ T cells are key mediators that control viral burden during chronic infection, we evaluated antigen-specific CD8^+^ T cells identified by tetramer staining for cells recognizing GP33, the major MHC Class Ib-restricted epitope of LCMV. Over the course of chronic infection, TCF-1^+^Tim3^-^ progenitor stem-like Tex cells (Tex^PROG^) maintain responses during infection and give rise to TCF-1^-^Tim3^+^ terminally exhausted T cells (Tex^TERM^) ^4^. However, expression of activated PI3Kδ led to a significant reduction in the percentage and numbers of splenic TCF-1^+^Tim3^-^Tex^PROG^ cells (**Fig.2A-B and Extended Data Fig.2A)**. Parallel findings were observed among antigen-specific CD8^+^ T cells in lung and liver of *Pik3cd*^E1020K/+^ mice, suggesting a broad loss of Tex^PROG^ cells (**Extended Data Fig.2B**). In contrast, no significant differences in percentages or numbers of TCF-1^-^Tim3^+^ Tex cells were observed, although Tim3 levels decreased over time (**Fig.2A-B and Extended Data Fig.2A**).

**Figure 2.**
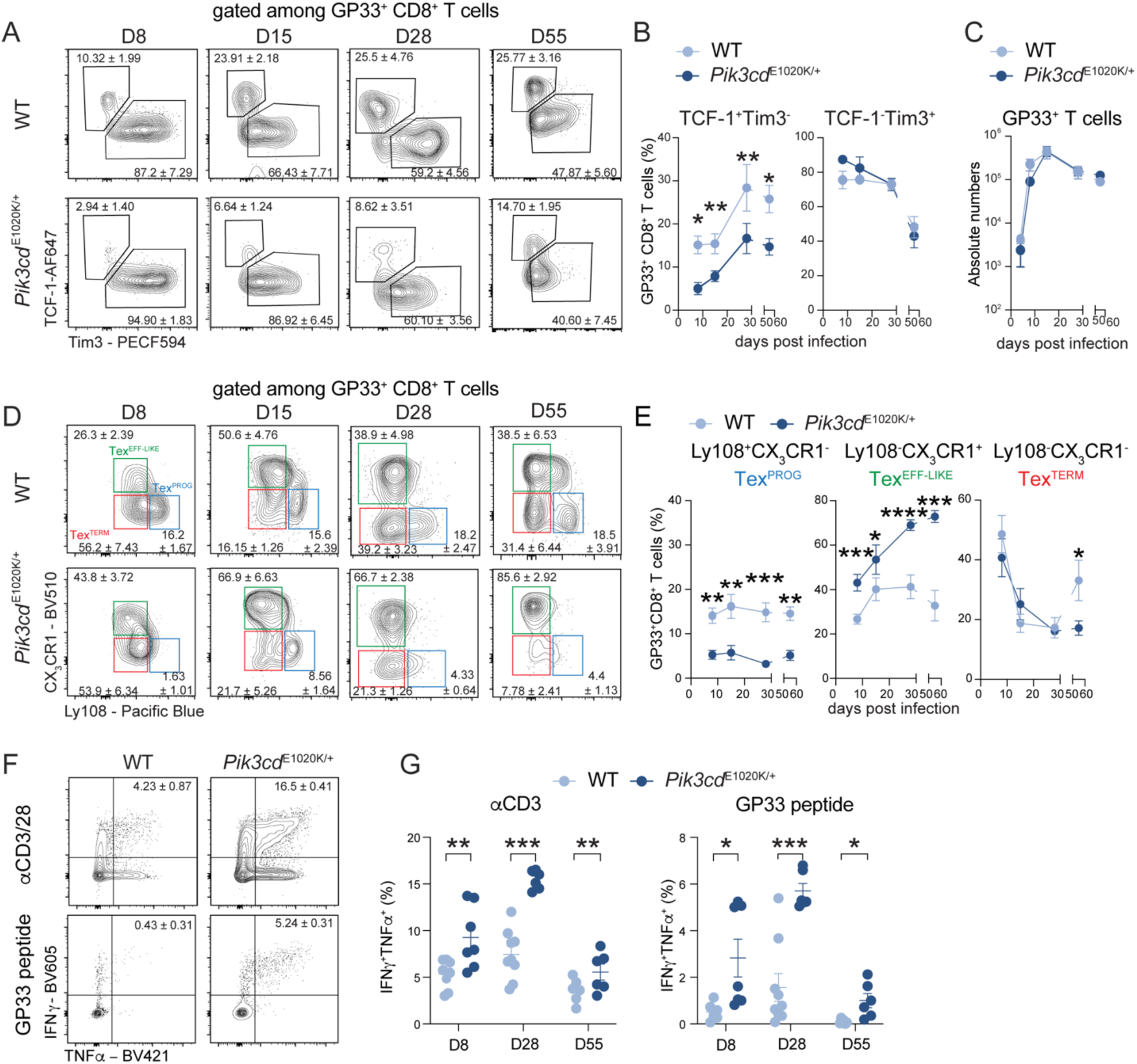
Activated PI3Kδ promotes expansion of CX_3_CR1^+^ effector-like Tex cells. C57BL/6 WT and *Pik3cd*^E1020K/+^ mice were infected i.v. with 2x10^6^ PFU of LCMV cl13. **(A-B)** Representative FACS plots **(A)** and graphs **(B)** of TCF-1^+^ Tim3^-^ and TCF-1^-^ Tim3^+^ cell percentages among GP33^+^ CD8^+^ T cells at indicated timepoints. **(C)** Absolute numbers of splenic GP33^+^ CD8^+^ T cells during chronic infection. **(D-E)** Representative FACS plots **(D)** and graphs **(E)** of Tex^PROG^, Tex^TERM^ and Tex^EFF-LIKE^ cell percentages among GP33^+^ CD8^+^ T cells. **(F-G)** Splenocytes from d28 p.i were stimulated with GP33 peptide or CD3/CD28 mAbs for 2 hrs. Representative FACS plots **(F)** and graphs **(G)** of IFNγ^+^TNFα^+^ CD8^+^ T cell percentage. Numbers in contour plots indicate means ± SEM of shown experiment. In **(B, C, E)** for WT condition, each dot represents pools of 11, 8, 8 and 10 mice at d8, d15, d28, d55 p.i. respectively and for *Pik3cd*^E1020K/+^ each dot represents pools of 11, 8, 7 and 10 mice, at d8, d15, d28, d55 p.i. respectively. In **(C)**, at day 4 time point, dots represent a pool of 4 WT and 3 *Pik3cd*^E1020K/+^ mice. In **(G)** each dot represents one mouse. Statistical differences were determined by one-way ANOVA **(B, C, E)** and Student’s t-test **(G)** *p < 0.05, **p < 0.01, ***p < 0.001, ****p <0.0001.

Decreased numbers of Tex^PROG^ cells are associated with a progressive loss of antigen-specific cells and decreased viral control during chronic infection ^3, 4, 5, 6, 7^. However, numbers and percentages of antigen-specific CD8^+^ T cells were similar between *Pik3cd*^E1020K/+^ mice and WT littermates in the spleen, lung and liver (**Fig.2C and Extended Data Fig.2C-D**). Activated PI3Kδ, therefore, did not affect T cell persistence, despite the decrease in Tex^PROG^ cells, suggesting alternative mechanisms contribute to the maintenance of antigen-specific cells in *Pik3cd*^E1020K/+^ mice during chronic infection.

Recent data suggest that during chronic infection, T cell populations are more heterogeneous than initially appreciated, with an effector-like (Tex^EFF-LIKE^) population that expresses the chemokine receptor CX_3_CR1, as well as KLRG1, T-bet and Zeb2. These CX_3_CR1^+^ cells persist during chronic infection and maintain effector functions ^38, 39^. Evaluation of cells using CX_3_CR1 and Ly108 (a marker for TCF-1^+^ Tex^PROG^ cells) ^40^, confirmed decreased CX_3_CR1^-^Ly108^+^ Tex^PROG^ in *Pik3cd*^E1020K/+^ mice with minimal changes in CX_3_CR1^-^Ly108^-^ terminally exhausted (Tex^TERM^) cells compared to WT **(Fig.2D-E and Extended Data Fig.2E).** In contrast, *Pik3cd*^E1020K/+^ mice displayed a large expansion of the Tex^EFF-LIKE^ population in spleen **(Fig.2D-E)**, lung and liver **(Extended Data Fig.2F-G)** compared to WT controls. Moreover, CD8^+^ T cells from *Pik3cd*^E1020K/+^mice displayed and maintained increased capacity to produce pro-inflammatory cytokines TNFα and IFNγ upon *ex vivo* peptide or CD3/28 stimulation (**Fig.2F-G**). These observations suggest that activated PI3Kδ drives a shift to effector-like CD8^+^ T cells, with enhanced persistence and function that may contribute to both immunedysregulation and the improved viral control during chronic infection.

### Loss of Tex^PROG^ cells in *Pik3cd*^E1020K/+^ mice is FoxO1-dependent

To assess if activated PI3Kδ drives these phenotypes through CD8^+^ T cell-intrinsic mechanisms, antigen-specific WT CD45.1 and *Pik3cd*^E1020K/+^ CD45.2 P14 T cell receptor transgenic cells were co-transferred into CD45.1/2 hosts that were subsequently infected with LCMV cl13 (**Fig.3A)**. Under these conditions, we did not observe increased mortality **(Extended Data Fig.3A)** and the impact of activated PI3Kδ P14 CD8^+^ T cells could be compared to WT P14 cells in the same environment, on a fixed TCR background, exposed to the same viral load. Although transfers of WT or *Pik3cd*^E1020K/+^ P14 cells individually showed no significant differences in cell numbers at d8 p.i. (**Extended Data Fig.3B**), *Pik3cd*^E1020K/+^ P14 cells did not persist as well as their WT counterparts in the competitive setting (**Fig.3B and Extended Data Fig.3C**). Furthermore, in both types of transfers, including the co-transfers where WT and mutant cells were present in the same mice, *Pik3cd*^E1020K/+^ P14 cells exhibited decreased percentages of Tex^PROG^ cells compared to WT (co-transfers shown in **Fig.3C**). *Pik3cd*^E1020K/+^ P14 cells also displayed decreased levels of TOX, which is critical for the maintenance of Tex cells, compared to WT control cells ^41^ (**Fig.3D).** Finally, percentages of the CX_3_CR1^+^ Tex^EFF-LIKE^ subset were significantly increased in *Pik3cd*^E1020K/+^ compared to WT P14 donors (**Fig.3C)**, providing evidence for CD8^+^ T cell-intrinsic contributions to the altered differentiation program of *Pik3cd*^E1020K/+^ CD8^+^ T cells.

**Figure 3.**
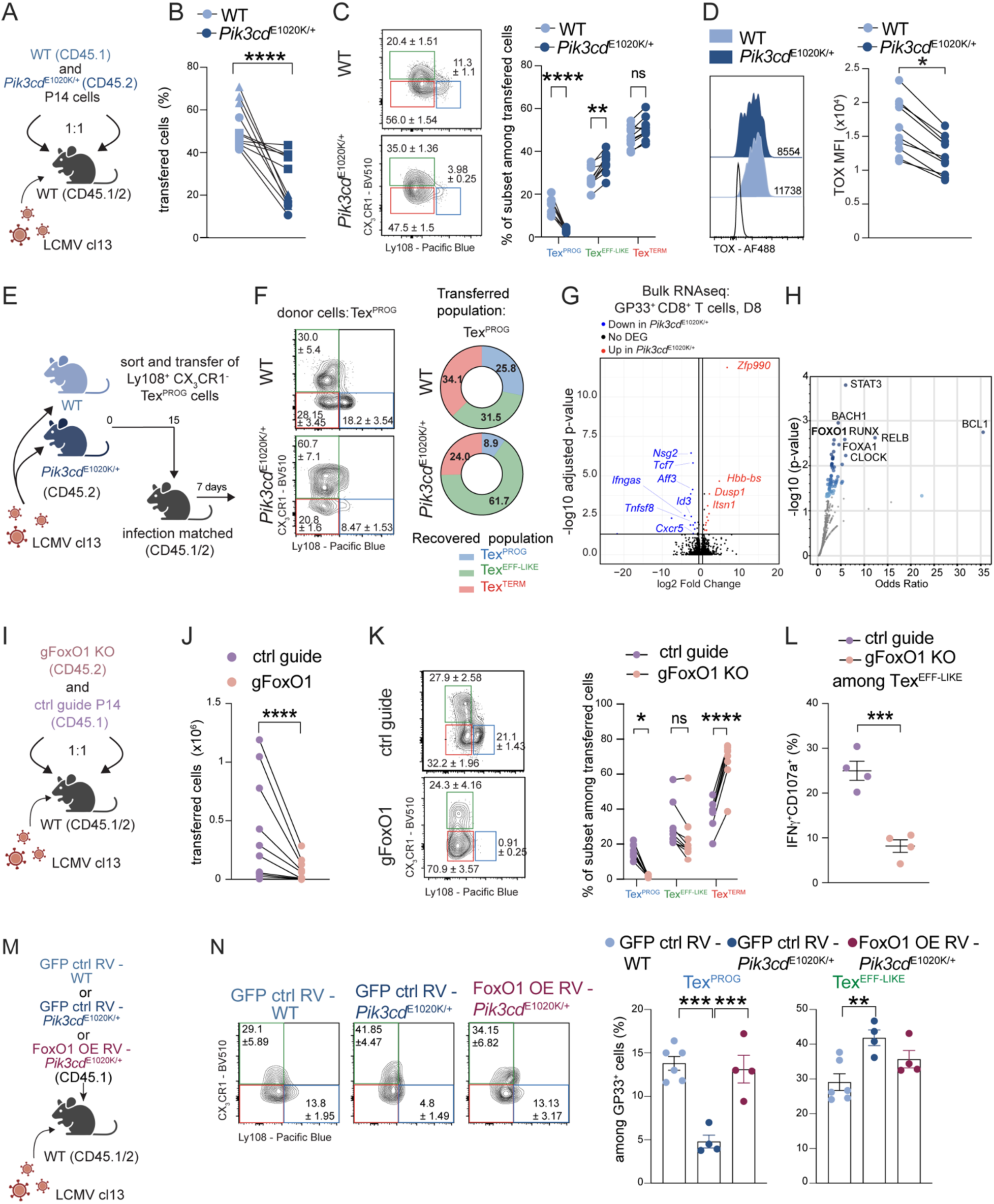
Loss of Tex^PROG^ in activated PI3Kδ mice is FoxO1 dependent. **(A-C)** CD8^+^ T cells sorted from C57BL/6 P14 (CD45.1) mice and *Pik3cd*^E1020K/+^ P14 (CD45.2) mice were co-transferred (2.5x10^3^ cells from each genotype) into congenic C57BL/6 (CD45.1/2) mice, which were infected i.v. with 2x10^6^ PFU of LCMV cl13. Transferred cells were analyzed d8 p.i.. **(A)** Experimental design. **(B)** Percentages of recovered transferred P14 cells. Each symbol shape represents one experiment. **(C)** Representative FACS plots (left) and graphs (right) of indicated population. **(D)** Representative histogram (left) and graph (right) of TOX MFI in P14 cells. **(E-F)** C57BL/6 WT and *Pik3cd*^E1020K/+^ mice were infected with LCMV cl13. **(E-F)** D15 p.i. sorted Tex^PROG^ cells (50-100x10^3^) from C57BL/6 WT or *Pik3cd*^E1020K/+^ infected mice were transferred into infection matched host. Transferred cells were analyzed d7 post-transfer. **(E)** Experimental design. **(F)** Representative FACS plots (left) and donut plots (right) showing percentages of indicated recovered population. **(G-H)** GP33^+^ CD8^+^ T cells d8 p.i. were sorted and analyzed by bulk RNA sequencing. **(G)** Volcano plot of DEGs, with significantly enriched gene sets colored in blue or red (FC < 1.5, p<0.05). **(H)** Pathway enrichment of TF Perturbations Followed by Expression gene sets, with significantly enriched gene sets indicated in blue. FoxO1 indicated in bold. **(I-L)** CD45.1 CRISPR-Cas9 control and CD45.2 CRISPR-Cas9 FoxO1-deficient P14 CD8^+^ T cells were co-transferred into congenic C57BL/6 (CD45.1/2) mice, which were infected with LCMV cl13. Transferred cells were analyzed d8 p.i.. **(I)** Experimental design. **(J)** Absolute numbers of transferred cells on d8 p.i.. **(K)** Representative FACS plots (left) and graph (right) of percentages of indicated population among transferred cells. **(L)** Percentages of IFNγ^+^CD107a^+^ after *ex vivo* GP33 peptide stimulation among Tex^EFF-LIKE^ cells. **(M-N)** Retroviral (RV)-mediated overexpression (OE) of FoxO1 in *Pik3cd*^E1020K/+^ (FoxO1 OE RV-*Pik3cd*^E1020K/+^), control WT GFP-RV (GFP ctrl RV-WT) or control *Pik3cd*^E1020K/+^ GFP-RV (GFP ctrl RV-*Pik3cd*^E1020K/+^) P14 cells (5x10^3^) were transferred into WT hosts and infected with LCMV cl13. And cells were analyzed d8 p.i.. **(M)** Experimental design. **(N)** Representative FACS plots (left) and graphs plot (right) of indicated population. Data are a pool of at least two independent experiments, with at least 3 mice per group. In **(L)**, data are representative of two independent experiments, with 4-5 mice per group. Numbers in contour plots indicate means ± SEM of shown experiment. Donut plot showing means of 3 pooled experiments. In **(B-D), (J-L), (N),** each dot represents 1 mouse. Statistical differences were determined by paired two-tailed *t* tests **(B, C, D, J, K)** and Student’s t-test **(L, N)** *p< 0.05, **p < 0.01, ***p < 0.001, ****p <0.0001.

To evaluate the developmental potential of Tex^PROG^ cells, equivalent numbers of sorted polyclonal Ly108^+^CX_3_CR1^-^ Tex^PROG^ cells from d15 LCMV cl13 infected *Pik3cd*^E1020K/+^ or WT CD45.2 donors were transferred into infection-matched CD45.1/2 hosts and assessed 1 week later (**Fig.3E and Extended Data Fig.3D**). Polyclonal Tex^PROG^ cells were used due to the low numbers of GP33^+^ or P14 antigen-specific *Pik3cd*^E1020K/+^ Tex^PROG^ cells obtained post-infection. Equivalent numbers of cells were maintained when polyclonal Tex^PROG^ cells were transferred from either genotype, confirming that *Pik3cd*^E1020K/+^ cells could persist under conditions inducing exhaustion **(Extended Data Fig.3E).** However, fewer Tex^PROG^ cells (9 **±** 1.53%) from *Pik3cd*^E1020K/+^ mice remained as Tex^PROG^ cells compared to those from WT Tex^PROG^ (25.8 **±** 7.1%). Instead, *Pik3cd*^E1020K/+^ Tex^PROG^ cells preferentially differentiated into CX_3_CR1^+^ Tex^EFF-LIKE^ cells (61.7 **±** 3.7%), whereas WT Tex^PROG^ differentiated similarly to either Tex^TERM^ (34.2+/- 7.9 %) or Tex^EFF-LIKE^ (31.5 **±** 4.7%) subsets (**Fig.3F and Extended Data Fig.3F**). Thus, activated PI3Kδ impairs the maintenance of CD8^+^ Tex^PROG^ cells, accelerating their transition towards a Tex^EFF-LIKE^ cell fate during chronic infection.

To define mechanism(s) by which activated PI3Kδ alters Tex cell differentiation, we evaluated transcriptomes of antigen-specific CD8^+^ T cells. Bulk RNA sequencing (RNAseq) on sorted GP33^+^ CD8^+^ T cells on d8 p.i. from *Pik3cd*^E1020K/+^ and WT littermates revealed only 25 differentially expressed genes (DEG) with greater than 1.5-fold differences between WT and *Pik3cd*^E1020K/+^ cells **(Fig.3G, Supplementary Table 1)**; many of these DEGs were key signature genes associated with Tex^PROG^ cells such as *Tcf7* (encoding TCF-1)*, Aff3, Id3, Nsg2* and *Cxcr5*, consistent with the reduction of this cell population. TF perturbation analysis (using ChIP-X Enrichment Analysis 3 ^42^) indicated that one of the top perturbated TF targets in *Pik3cd*^E1020K/+^ cells compared to WT was the PI3K-regulated TF FoxO1, which is required for TCF-1 expression (**Fig.3H, Supplementary Table 2**).

PI3Kδ activation induces the activation of Akt, which phosphorylates FoxO1, leading to FoxO1 exclusion from the nucleus and targeted degradation in the cytosol (**Extended Data Fig.3G**) ^43^. To address the role of FoxO1 in Tex cell subset differentiation, we used CRISPR-Cas9 ribonucleoproteins (RNPs) to disrupt *Foxo1* expression in WT P14 cells. Control-RNP and *FoxO1*-RNP treated P14 cells were co-transferred into CD45.1/2 mice, which were infected with LCMV cl13 and analyzed 7 days later (**Fig.3I**). FoxO1-depletion as well as the loss of TCF-1 were validated by flow cytometry (**Extended Data Fig.3H-I**). FoxO1-deficient P14 CD8^+^ T cells did not persist as well as WT cells in the same host (**Fig.3J**), similar to previous reports ^23, 25^. FoxO1 depletion also led to the loss of Tex^PROG^ cells, as previously reported ^23, 25^ (**Fig.3K)**. However, FoxO1 depletion did not increase CX_3_CR1^+^ Tex^EFF-LIKE^ cells or effector function, although KLRG1 expression was increased (**Fig.3K-L and Extended Data Fig.3J**). FoxO1 depletion also decreased TOX levels, but did not affect inhibitory receptor expression on P14 cells (**Extended Data Fig.3K-L**), despite previous data showing that FoxO1 was required to maintain PD-1 expression during exhaustion ^23^. Conversely, retroviral-mediated overexpression of FoxO1 in *Pik3cd*^E1020K/+^ P14 cells followed by transfer into WT hosts, rescued Tex^PROG^ cell percentages, but only partially reduced the expansion of Tex^EFF-LIKE^ cells (**Fig.3M-N**). These results implicate the disruption of FoxO1 in the cell-intrinsic loss of *Pik3cd*^E1020K/+^ Tex^PROG^ cells, but suggest that other factors influence the increase in Tex^EFF-LIKE^ cells.

### scRNAseq reveals early PI3Kδ mediated reprogramming of Tex^PROG^ towards effector fate

To define molecular mechanisms by which activated PI3Kδ alters Tex^PROG^ cell differentiation, we performed single cell RNA sequencing (scRNAseq) at an early time-point during infection (d8 p.i.) on sorted antigen-specific (GP33^+^) CD8^+^ T cells. Among the 7 clusters identified (**Fig.4A-B**), the Tex^PROG^ cell cluster (defined by *Slamf6* encoding Ly108*, Id3*, *Tcf7*, and *Tox)* was reduced in *Pik3cd*^E1020K/+^ cells compared to WT (**Fig.4A-C)**, analogous to the phenotype observed by flow cytometry (see **Fig.2D**). Intriguingly, *Pik3cd*^E1020K/+^ Tex^PROG^ cells displayed both a reduced expression of genes associated with memory and stemness (e.g. *Id3, Myb* and *Il7r)* **(Fig.4D, Supplementary Table 3)** ^44, 45^ and increased expression of genes associated with effector cells (e.g. *Zeb2*, *Prdm1, Id2* and *Tbx21)* compared to WT. Thus, PI3Kδ activation in the early phase of chronic infection shifts the Tex^PROG^ stemness program toward an effector-like profile.

**Figure 4.**
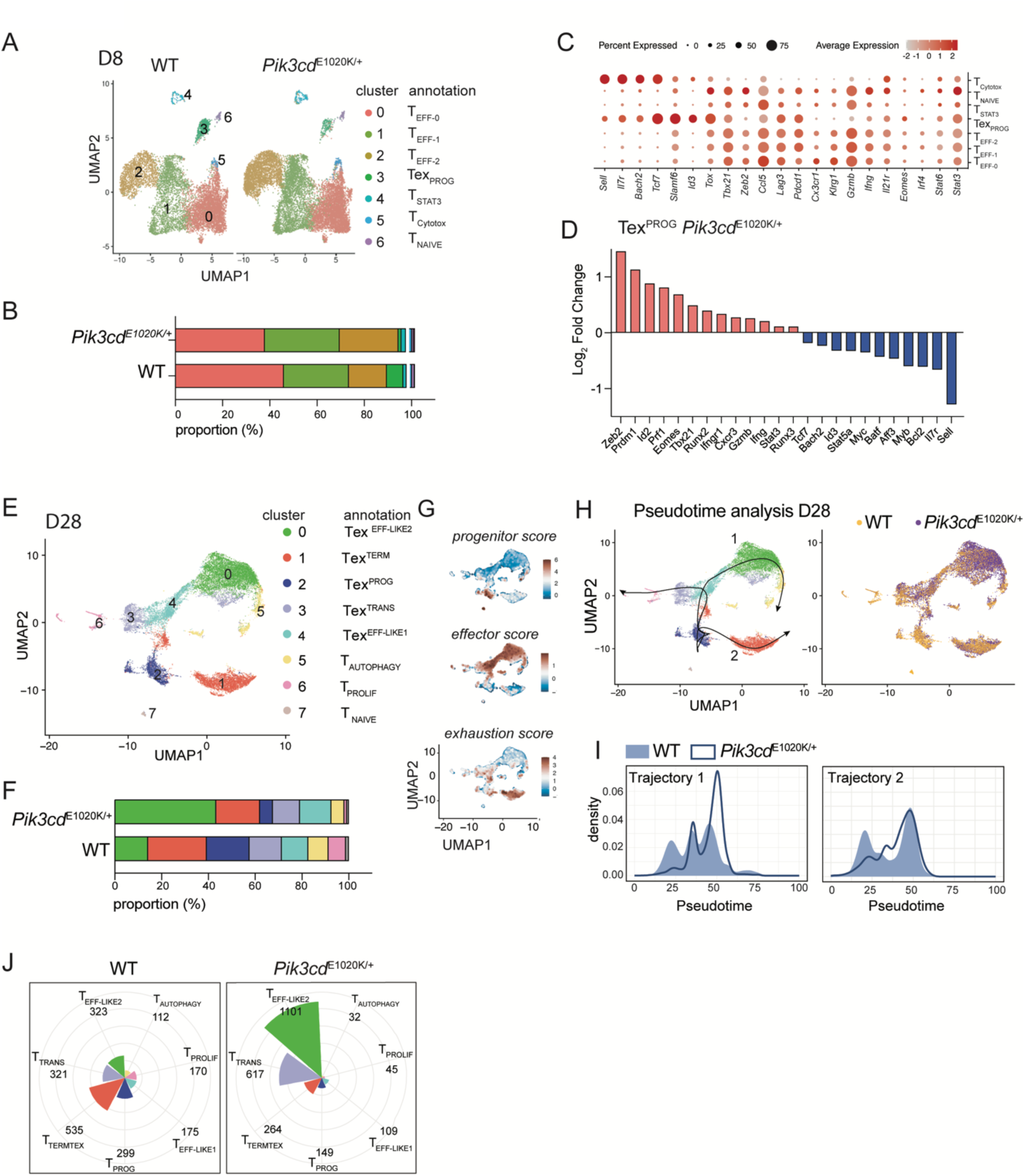
scRNAseq reveals early Tex^EFF-LIKE^ differentiation and expansion in activated PI3Kδ mice. C57BL/6 WT and *Pik3cd*^E1020K/+^ mice were infected i.v. with 2x10^6^ PFU of LCMV cl13 and GP33^+^ CD8^+^ T cells were sorted and analyzed on d8 **(A-D)** and d28 p.i. **(E-K)** for single cell RNAseq. **(A)** Uniform manifold approximation and mapping (UMAP) plotting of RNA-defined Seurat clusters from GP33^+^ CD8^+^ T cells d8 p.i.. **(B)** Bar plot of cells per cluster among WT and *Pik3cd*^E1020K/+^ mice. **(C)** Dot plot showing expression of indicated gene in each cluster. Color scale indicates *Z* score of gene expression. **(D)** Bar plot showing fold change of selected progenitor and effector genes among Tex^PROG^ genes *Pik3cd*^E1020K/+^ compared to WT. **(E)** UMAP plotting of RNA-defined Seurat clusters from GP33^+^ CD8^+^ T cells d28 p.i.. **(F)** Bar plot of cells per clusters among WT and *Pik3cd*^E1020K/+^ mice. **(G)** Feature plots showing indicated module score**. (H)** Slingshot trajectory analysis projected on UMAP showing main trajectories (left) and UMAP highlighting WT and *Pik3cd*^E1020K/+^ cells (right). **(I)** Histogram showing cell density of WT and *Pik3cd*^E1020K/+^ cells on indicated trajectory. **(J)** Rose chart comparing number of cells in each cluster from top 10 clonotypes in WT and *Pik3cd*^E1020K/+^ mice. The axis scale is consistent across both charts.

### PI3Kδ promotes differentiation and expansion of Tex^EFF-LIKE^

Because the loss of FoxO1 did not recapitulate the effector expansion in *Pik3cd*^E1020K/+^ CD8^+^ T cells (see **Fig.3K and N**), we reasoned that other factors contribute to this phenotype. To further characterize the consequences on PI3Kδ activation of Tex^EFF-LIKE^ generation, we analyzed antigen-specific (GP33^+^) CD8^+^ T cells by scRNAseq in both early (d8 p.i.) and late (d28 p.i.) phases of the infection. In the early phase of infection, *Pik3cd*^E1020K/+^ GP33^+^ CD8^+^ T cells showed an expansion of two effector-like clusters (1 and 2); these clusters were enriched for gene signatures related to cell division (e.g. cell cycle and mitotic spindle organization) compared to WT **(Extended Data Fig.4A, Supplementary Table 4)**, suggesting that activated PI3Kδ leads to an increase in activated effector CD8^+^ T cells with proliferative signatures.

At d28, 8 clusters were identified that were grouped into 3 phenotypes defined by key signature genes and modules as progenitor (cluster 2), effector (clusters 4 and 0, denoted Tex^EFF-LIKE1^ and Tex^EFF-LIKE2^, respectively) and exhausted cells (cluster 1) **(Fig.4E-G and Extended Data Fig.4B-C, Supplementary Table 5)**. Again, *Pik3cd*^E1020K/+^ GP33^+^ CD8^+^ T cells showed a decreased Tex^PROG^ population with increased representation of the Tex^EFF-LIKE^ clusters, especially Tex^EFF-LIKE2^ cells. The Tex^EFF-LIKE1^ cluster shared features with previously described “intermediate” Tex cells, such as increased CXCR6 expression compared to Tex^EFF-LIKE2^ cells ^40^, although multiple effector-associated genes, including *Tbx21* and *Zeb2,* were increased compared to Tex^EFF-LIKE2^ cells (**Extended Data Fig.4B-C**).

Consistent with early skewing towards an effector fate in *Pik3cd*^E1020K/+^ cells, Slingshot trajectory analysis on d28 p.i. revealed that Tex^PROG^ from *Pik3cd*^E1020K/+^ mice were preferentially found on an effector-like trajectory (trajectory 1), while cells of both WT and *Pik3cd*^E1020K/+^ genotypes appeared to commit similarly to an exhaustion path (trajectory 2) **(Fig.4H-I)**. In addition, single cell TCR sequencing and evaluation of the top 10 expanded T cell clones revealed that major clonal expansions occurred primarily in Tex^EFF-LIKE2^ cells from *Pik3cd*^E1020K/+^ mice, whereas WT mice showed diverse patterns of clonal expansion with increased Tex^TERM^ clones at d28 p.i. (**Fig.4J**). In conjunction with the increased cell-cycle signatures observed in T^EFF^ clusters on d8 p.i., these data suggest that activated PI3Kδ promotes both the differentiation and expansion of effector cells under conditions that induce exhaustion.

### CX_3_CR1^+^ Tex^EFF-LIKE^ cell expansion and persistence is CD4 and IL-21 dependent

Pathway analysis comparing DEG from pseudobulk aggregated antigen-specific CD8^+^ T cells from WT and *Pik3cd*^E1020K/+^ mice on d28 p.i. revealed that IFNα and IFNγ response pathways were enriched specifically in WT cells, while signatures associated with IL2-STAT5 pathways were upregulated in both genotypes **(Fig.5A, Supplementary Table 6)**. In contrast, activated PI3Kδ CD8^+^ T cells showed increased induction of PI3K-AKT-mTOR pathways, as expected, but also a JAK-STAT3-IL6 signature **(Fig.5A)**. Analyses of DEGs affected by activated PI3Kδ on d8 p.i. CD8^+^ T cells bulk RNAseq further indicated that STAT3 targets constituted one of the most altered set of transcriptional targets in these cells (see **Fig.3H**). These results suggest that the skewing of Tex^PROG^ towards effector phenotypes may occur via differential activation of STAT proteins in the presence of activated PI3Kδ.

**Figure 5.**
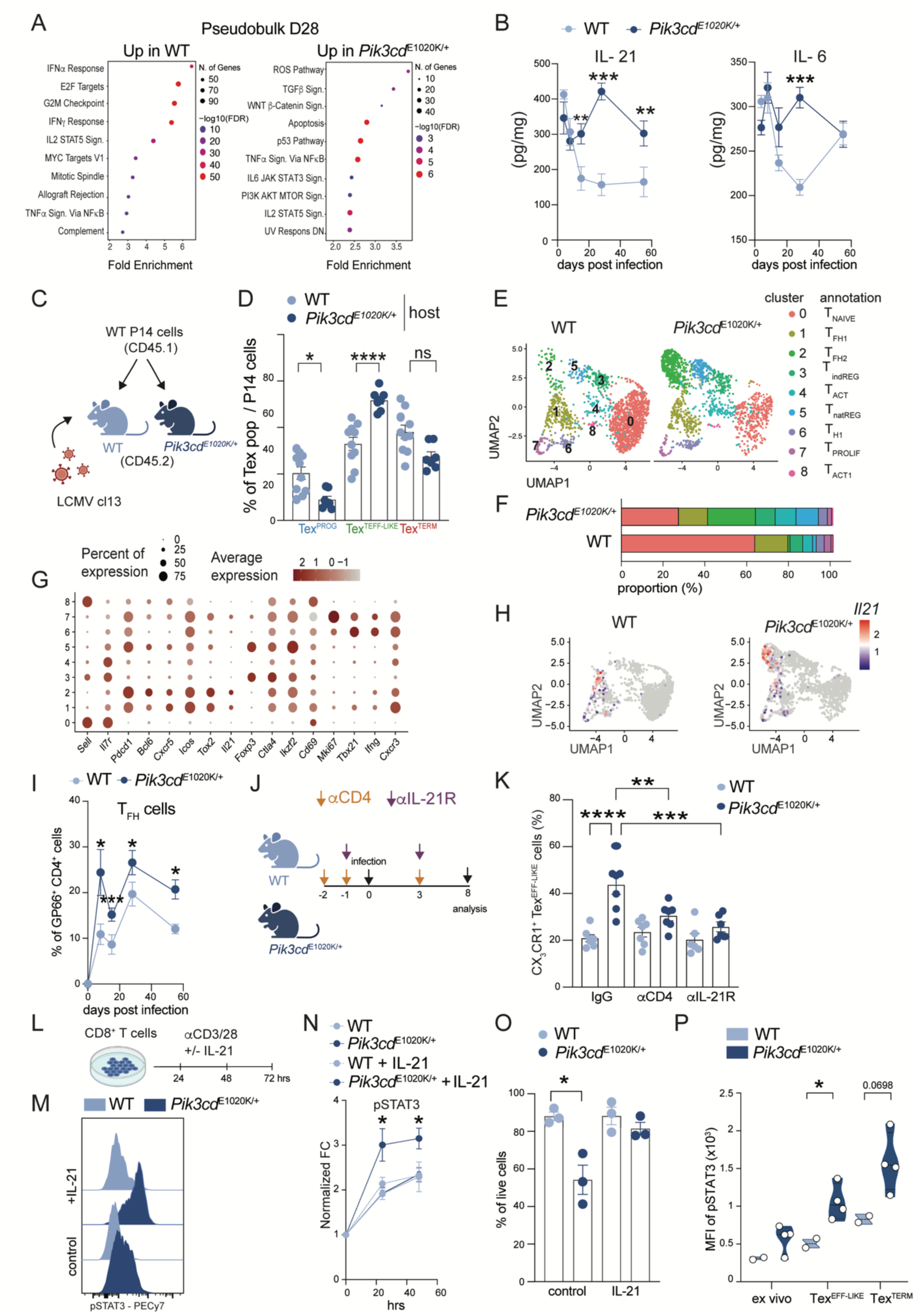
CX_3_CR1^+^ Tex^EFF-LIKE^ cell expansion and persistence is CD4 and IL-21 dependent. **(A)** Pseudobulk analysis of d28 p.i. scRNAseq data, showing HALLMARK pathway enrichment analysis of top DEGs between C57BL/6 WT and *Pik3cd*^E1020K/+^ mice. **(B)** IL-21 and IL-6 in liver homogenates at indicated timepoints. **(C-D)** CD8^+^ T cells were sorted from C57BL/6 P14 (CD45.1) mice and 5x10^3^ cells were transferred into WT or *Pik3cd*^E1020K/+^ CD45.2 mice, which were infected with LCMV cl13. **(C)** Experimental design. **(D)** Percentages of Tex^PROG^, Tex^EFF-LIKE^ and Tex^TERM^ cells in different genotype hosts at d8 p.i.. **(E-H)** C57BL/6 WT and *Pik3cd*^E1020K/+^ mice were infected i.v. with LCMV cl13. CD4^+^ T cells were sorted at d8 p.i. from WT C57BL/6 and *Pik3cd*^E1020K/+^ mice and analyzed by scRNAseq. **(E)** UMAP plotting of RNA-defined Seurat clusters of CD4^+^ T cells d8 p.i.. **(F)** Bar plot of cells per cluster among WT and *Pik3cd*^E1020K/+^ mice. **(G)** Dot plot showing expression level of indicated genes in each cluster. Color scale indicates *Z* score of gene expression. **(H)** Feature plot of *Il21* expression. **(I)** T_FH_ cells among GP66^+^ CD4^+^ T cells at indicated timepoints. **(J-K)** C57BL/6 or *Pik3cd*^E1020K/+^ mice were infected with LCMV cl13. Mice were either treated with depleting anti-CD4 or blocking IL-21R mAbs. Antigen-specific cells were analyzed d8 p.i.. **(J)** Experimental design. **(K)** Percentages of CX_3_CR1^+^ Tex^EFF-LIKE^ cells. (**L-O**) CD8^+^ T cells were stimulated *in vitro* for 3 days with anti-CD3/28 mAbs in the presence or absence of IL-21 and pSTAT3 and viability analyzed by flow cytometry. **(L)** Experimental design. **(M-N)** Representative pSTAT3 histogram **(M)** and graph **(N)** showing normalized fold change of pSTAT3. **(O)** Percentages of live cells after *in vitro* IL-21 stimulation. **(P)** CD8^+^ T cells isolated from d21 LCMV cl13 infected C57BL/6 WT and *Pik3cd*^E1020K/+^ mice were stimulated for 2 hrs *ex vivo* with IL-21 and analyzed by flow cytometry. Graph showing MFI of pSTAT3. Data are a pool of at least two independent experiments of at least 3 mice per group. In **(D, K and P)** each dot represents one mouse. In **(B)** in WT condition, each dot represents a pool of 4, 8, 4, 8, 4 mice at d4, d8, d15, d28 and d55 p.i. respectively and in *Pik3cd*^E1020K/+^ condition, each dot represents a pool of 4, 7, 4, 8, 4 mice at d4, d8, d15, d28 and d55. In **(I)** in WT condition, each dot represents a pool of 6, 6, 7, 7, 9 mice at d4, d8, d15, d28 and d55 p.i. respectively and in *Pik3cd*^E1020K/+^ condition, each dot represents a pool of 6, 7, 6, 7, 9 mice at d4, d8, d15, d28 and d55. In **(N)** one dot represent the mean of 3 experiment. In **(O)** one dot represents the mean of one experiment. Statistical differences were determined by Student’s t-test **(O)** and by one-way ANOVA **(B, D, I, K, N, P)** *p < 0.05, **p < 0.01, ***p< 0.001, ****p <0.0001.

STAT3 is a member of the signal transducer and activators of transcription (STAT) family, which are phosphorylated and activated by diverse cytokines ^46^. The increased induction of STAT3 pathways in CD8^+^ T cells from *Pik3cd*^E1020K/+^ mice suggested that an altered cytokine environment or cytokine responsiveness may contribute to their phenotypes. Indeed, multiple inflammatory cytokines were increased in liver homogenates from infected *Pik3cd*^E1020K/+^ mice relative to WT by 2 weeks p.i., including IL-6 and IL-21, two cytokines that activate STAT3 (**Fig.5B and Extended Data Fig.5A**) ^47^. IL-21 has also been shown to be important for responses to chronic viral infection ^48, 49^.

Although we observed cell-intrinsic effects of activated PI3Kδ on CD8^+^ T cell differentiation, the altered cytokine patterns in *Pik3cd*^E1020K/+^ mice suggested there may also be CD8^+^ T cell-extrinsic contributions. To evaluate this question, WT CD45.1 P14 cells were transferred into co-housed CD45.2 *Pik3cd*^E1020K/+^ or WT littermates, which were then infected with LCMV cl13 and analyzed at d8 (**Fig.5C**). We observed decreased Tex^PROG^ cell percentages among the transferred WT P14 cells in *Pik3cd*^E1020K/+^ hosts (**Fig.5D**). Moreover, WT P14 cells were strongly skewed towards CX_3_CR1^+^ Tex^EFF-LIKE^ population when transferred into *Pik3cd*^E1020K/+^ compared to WT hosts (**Fig.5D**). These results suggest that extrinsic factors contribute to both Tex^PROG^ cell persistence and expansion of Tex^EFF-LIKE^ in activated PI3Kδ mice as compared to WT.

Recent data indicate that IL-21 produced by T follicular helper cells (T_FH_) cells promotes Tex^EFF-LIKE^ antigen-specific Tex cells in the context of chronic infection ^49^. To evaluate CD4^+^ T cells in response to LCMV cl13, we performed scRNAseq on total CD4^+^ T cells from WT and *Pik3cd*^E1020K/+^ mice on d8 p.i.. By gene expression, 9 clusters were identified (**Fig.5E-F**), including naïve, T_FH_, T_REG_ and T_H1_ cells, defined by key signature genes **(Extended Data Fig.5B)**. Among these, two major clusters expressing T_FH_-associated genes (*Cxcr5*, *Pcdc1, Icos and Bcl6)* were identified: cluster 1 had higher *Cxcr3,* and lower *Pdcd1* and *Bcl6*, and was present in both genotypes **(Fig.5G and Extended Data Fig.5B)**, whereas Cluster 2 had higher levels of *Bcl6* and *Pdcd1* and was greatly expanded in *Pik3cd*^E1020K/+^ mice (**Fig.5E-G)**, consistent with our previous findings of increased germinal centers and T_FH_ in these mice [34]. Accordingly, we found increased percentages and numbers of GP_66-77_-antigen-specific CXCR5^+^PD-1^+^ T_FH_ cells in *Pik3cd*^E1020K/+^ compared to WT mice during chronic infection, although numbers of total T_FH_ post-infection did not differ (**Fig.5H-I and Extended Data Fig.5C-D)**. *Pik3cd*^E1020K/+^ mice also showed expansion of an activated T_REG_ population (cluster 5) similar to those described in mice expressing *Pik3cd*^E1020K/+^ specifically in T_REGS_, which also have increased T_FH_ cells ^50^. Notably, IL-21 was primarily expressed by the T_FH_ clusters, and was prominent in cluster 2, the T_FH_ cluster that was greatly expanded in *Pik3cd*^E1020K/+^ mice (**Fig.5H and Extended Data Fig.5E)**. Thus, *Pik3cd*^E1020K/+^ mice display increased percentages of a distinct *Il21*-expressing T_FH_ population during chronic infection.

To evaluate the effects of CD4^+^ T cells and IL-21 on phenotypes of *Pik3cd*^E1020K/+^ mice, we either depleted CD4^+^ T cells or blocked the IL-21R by injection of monoclonal antibodies **(Fig. 5J).** Mice were infected with LCMV cl13 and antigen-specific CD8^+^ T cells analyzed at d7; CD4 depletion was confirmed by flow cytometry (**Extended Data Fig.5F).** Depletion of CD4^+^ T cells or blocking IL-21R inhibited the expansion of CX_3_CR1^+^ Tex^EFF-LIKE^ cells specifically in *Pik3cd*^E1020K/+^ mice, yet did not significantly affect Tex^PROG^ or Tex^TERM^ cell percentages compared to control IgG injections, nor cell populations in WT mice at this early timepoint (**Fig.5K** and **Extended Data Fig.5G**). Thus, IL-21 is a key driver of the increased Tex^EFF-LIKE^ cells in response to LCMV cl13 in *Pik3cd*^E1020K/+^ mice.

### Activated PI3Kδ induces increased sensitivity to IL-21, promoting pSTAT3 and survival

The above data argue that an altered cytokine environment leads to the increased CD8^+^ effector T cells in *Pik3cd*^E1020K/+^ mice; however, we also observed a cell-intrinsic component to this phenotype. Because IL-21 signals via STAT3, we sought to determine the responsiveness of CD8^+^ T cell to IL-21. Sorted naïve CD8^+^ T cells were stimulated with anti-CD3/28 mAbs in the presence or absence of IL-21 for 3 days **(Fig.5L)**. IL-21 stimulation resulted in greater STAT3 phosphorylation (pSTAT3) in CD8^+^ T cells from *Pik3cd*^E1020K/+^ compared to WT mice **(Fig.5M-N**), despite similar levels of STAT3 protein **(Extended Data Fig.5H)**. TCR-mediated activation of *Pik3cd*^E1020K/+^ CD8^+^ T cells is associated with increased cell death ^27, 51^; however, IL-21 increased the survival of activated *Pik3cd*^E1020K/+^, but not WT CD8^+^ T cells, suggesting that IL-21 affects both their survival and differentiation (**Fig.5O**). To directly assess IL-21 responsiveness, activated CD8^+^ T cells were rested in serum free media to reduce background levels of pSTAT3, and then stimulated with IL-21 (**Extended Data Fig.5I)**. Again, *Pik3cd*^E1020K/+^ CD8^+^ T cells displayed elevated and sustained pSTAT3 in response to IL-21 compared to that seen in WT cells (**Extended Data Fig.5J)**. Furthermore, pSTAT3 was increased in CX_3_CR1^+^ Tex^EFF-LIKE^ cells and Tex^TERM^ from d21 p.i. *Pik3cd*^E1020K/+^ mice compared to those from WT mice when stimulated *ex vivo* with IL-21 (**Fig.5P and Extended Data Fig.5K**). Consistent with these observations, expression of *Il21r* was increased, while *Socs1* and *3*, which encode two key inhibitors of STAT3 and JAK phosphorylation ^52^, were decreased in CD8^+^ T cells from *Pik3cd*^E1020K/+^ mice at d28 p.i. (**Extended Data Fig.5L-M**). These results provide support that both increased IL-21 production and responsiveness contribute to the expansion and persistence of *Pik3cd*^E1020K/+^ CX_3_CR1^+^ Tex^EFF-LIKE^ cells through heightened STAT3 phosphorylation and cell survival.

### Tex^TERM^ and Tex^PROG^ from *Pik3cd*^E1020K/+^ mice display decreased population identity

To better understand the consequences of activated PI3Kδ on CD8^+^ T cell differentiation during chronic infection, we further mined our RNAseq data set from d28 p.i.. Comparison of module scores revealed an increase in effector features in Tex^PROG^, Tex^EFF-LIKE1^ and Tex^TERM^ subsets from *Pik3cd*^E1020K/+^ mice **(Fig.6A, middle panel)**. Specific DEGs included increased expression of *FasL*, a key mediator of T cell apoptosis, in Tex^PROG^ cells (**Fig.6B, Supplementary Table 7)**; *Pik3cd*^E1020K/+^ Tex^EFF-LIKE1^ cells expressed higher levels of *Klrg1*, as well as mitochondrial and ribosomal genes (e.g. *Mt-Co2* and *Rpl28*), reflective of increased effector and synthetic function (**Fig.6B, Supplementary Table 7)**. Moreover, Tex^TERM^ cells from activated PI3Kδ mice also displayed higher levels of multiple effector-associated genes such as *Klrg1*, *Klrk1*, *Klrd1* and *Tnf* (**Fig.6B, Supplementary Table 7)**. Pathway analysis established that *Pik3cd*^E1020K/+^ Tex^TERM^ cells were enriched for ribosome and T cell signaling pathways suggesting these cells were more activated and synthetic, whereas WT Tex^TERM^ cells were enriched for cell-death associated pathways (**Extended Data Fig.6A**). However, *Pik3cd*^E1020K/+^ Tex^TERM^ cells also expressed higher levels of genes associated with stem-cell/memory populations, including *Lef1* and *IL7r* **(Fig.6B)** and were enriched for a progenitor signature **(Fig.6A, left panel)**.

**Figure 6.**
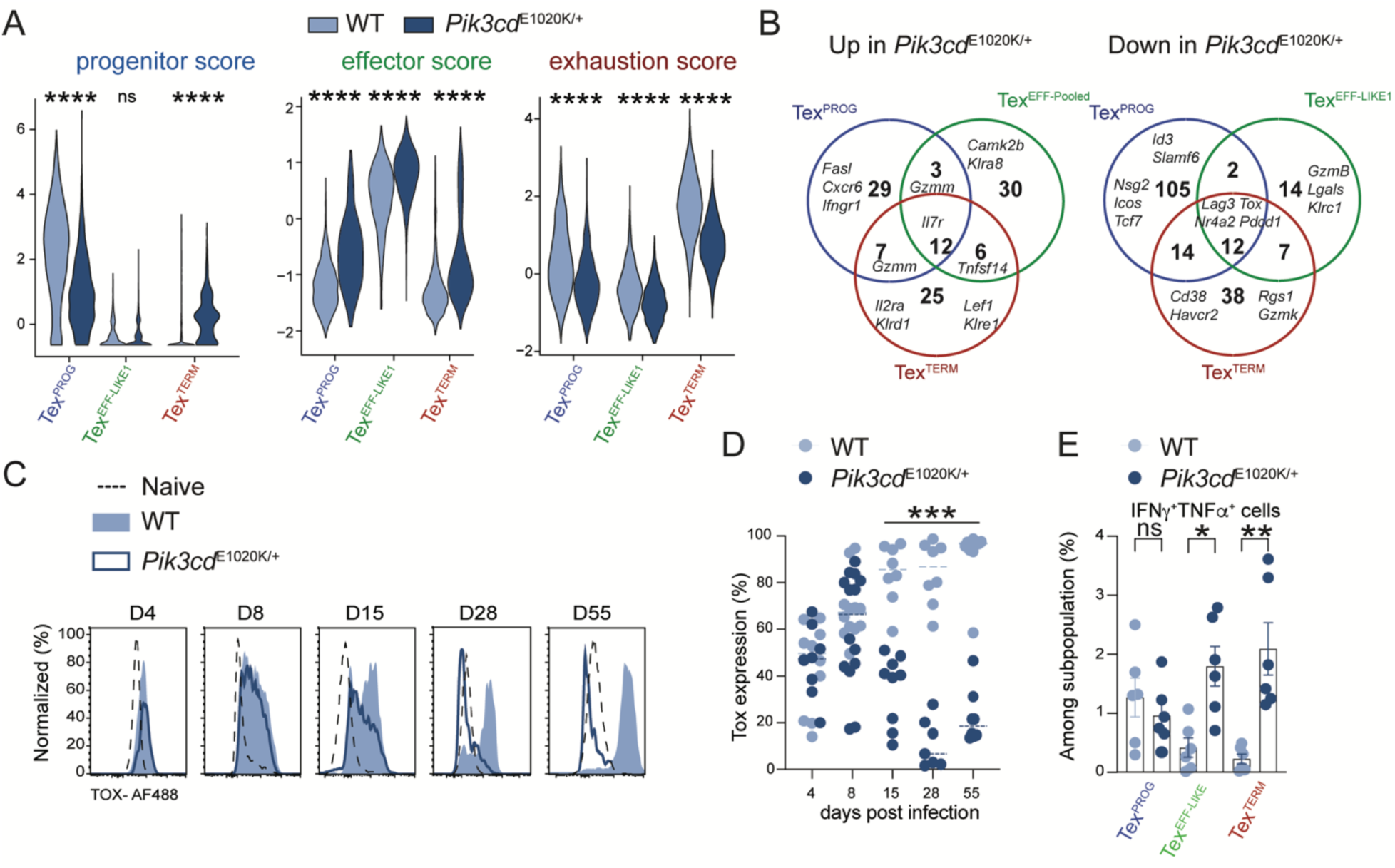
Activated PI3Kδ suppresses TOX and exhaustion program. **(A)** Violin plot of indicated module score among in WT and *Pik3cd*^E1020K/+^ Tex^PROG^, Tex^EFF-LIKE^ and Tex^TERM^ cells on d28 p.i.. **(B)** Venn diagram showing top upregulated DEGs (left) and downregulated (right) of indicated cell population from *Pik3cd*^E1020K/+^ relative to WT mice (p < 0.05). **(C-D)** Representative histograms **(C)** and graphs **(D)** of percentages of TOX at indicated timepoint. **(E)** Percentages of IFNγ^+^TNFα^+^ after *ex vivo* stimulation with GP33 peptide among indicated population at d28 p.i.. In **(D-E)** each point represents one mouse, data are a pool of at least two independent experiments of at least 3 mice per group. Statistical differences were determined by one-way ANOVA (**D**) and by Student’s t-test (**E)** *P < 0.05, **P < 0.01, ***P < 0.001.

In contrast, comparison of DEGs expressed at lower levels in *Pik3cd*^E1020K/+^ vs WT in Tex^PROG^ and Tex^TERM^ indicated a downregulation of key signature genes of these major clusters **(Fig.6A-B)**. For Tex^PROG^, this included reduced expression of *Id3* and *Slamf6 (***Fig.6B**), as well as a loss of *Myb*, which encodes a TF that is essential for an early progenitor population in exhaustion ^45^ (**Extended Data Fig.6B**). Tex^TERM^ showed decreased expression of *Havcr2* (encoding Tim3), while all 3 clusters exhibited decreased expression of exhaustion-associated genes *NR4a, Pdcd1, Lag3 and Tox* (**Fig.6B)**. These observations were confirmed at the protein level by flow cytometry, including reduced Tim3 and a near loss of LAG3, which plays a key role in repressing effector cell function during exhaustion ^53, 54, 55, 56^ (**Extended Data Fig.6C-D**). Notably, we observed significantly decreased expression of TOX in *Pik3cd*^E1020K/+^ antigen-specific CD8^+^ T cells starting at day 15 p.i. **(Fig.6C-D).** In contrast, KLRG1 was increased in both CX_3_CR1^+^ Tex^EFF-LIKE^ and a portion of Tex^TERM^ cells **(Extended Data Fig.6C-D)**. Accordingly, both CX_3_CR1^+^ Tex^EFF-LIKE^ and Tex^TERM^ CD8^+^ T cells from *Pik3cd*^E1020K/+^ mice displayed increased capacity to produce pro-inflammatory cytokines TNFα and IFNγ upon *ex vivo* peptide stimulation (**Fig.6E**). Notably, CD127 (IL-7Ra) expression was increased in *Pik3cd*^E1020K/+^ Tex^TERM^ cells **(Extended Data Fig.6C-D)**, consistent with the increased progenitor signature in these cells **(Fig.6A)**. The decrease of key signature markers, increased effector functions, as well as progenitor signature in Tex^TERM^ cells suggested an overall diminished commitment to the exhaustion program in the presence of activated PI3Kδ.

### Activated PI3Kδ prevents epigenetic features of exhaustion while promoting Tex^TERM^ plasticity

TOX plays a critical role in the establishment of epigenetic changes that help establish and maintain the exhaustion program in CD8^+^ T cells ^8, 9, 10, 11, 12, 41^. To investigate how activated PI3Kδ affects epigenetic features of exhaustion, Tex^PROG^ (Ly108^+^CX_3_R1^-^), Tex^EFF-LIKE^ (CX_3_CR1^+^Ly108^-^) and Tex^TERM^ (CX_3_CR1^-^Ly108^-^CD101^+^) antigen-specific CD8^+^ T cells were isolated at d15 p.i., a time point when exhaustion has been established, and used for assay for transposase-accessible chromatin using sequencing (ATACseq) (**Fig.7A**). CD101 was included as an additional marker to isolate terminally exhausted cells (see **Extended Data Fig.7B-C)** ^38^. Principal component analysis (PCA) of chromatin-accessible regions (ChARs) showed separation of *Pik3cd*^E1020K/+^ versus WT on PC2 with the different Tex subpopulations separating on PC1 (**Fig.7B**). Of note, *Pik3cd*^E1020K/+^ Tex^PROG^, Tex^EFF-LIKE^ and Tex^TERMTEX^ populations were shifted to the right relative to their WT equivalents on the PC1 axis, more towards the direction of Tex^EFF-LIKE^ cells. HOMER analyses ^57^ of differential ChaRs revealed that binding sites for several TF associated with effector function, including T-bet, Blimp1 (encoded by *Prdm1*, a target of STAT3 ^58^), IRF4 and BHLHE40 were enriched in *Pik3cd*^E1020K/+^ cell populations, particularly in Tex^PROG^ (**Fig.7C, Supplementary Table 8**), reflecting the increased effector module scores seen in gene expression (see **Fig.6A)**. In contrast, increased markings on STAT5 motifs were observed in all 3 Tex subpopulations from WT compared to activated PI3Kδ mice, confirming distinct features of WT and *Pik3cd*^E1020K/+^ cells (**Fig.7C**).

**Figure 7.**
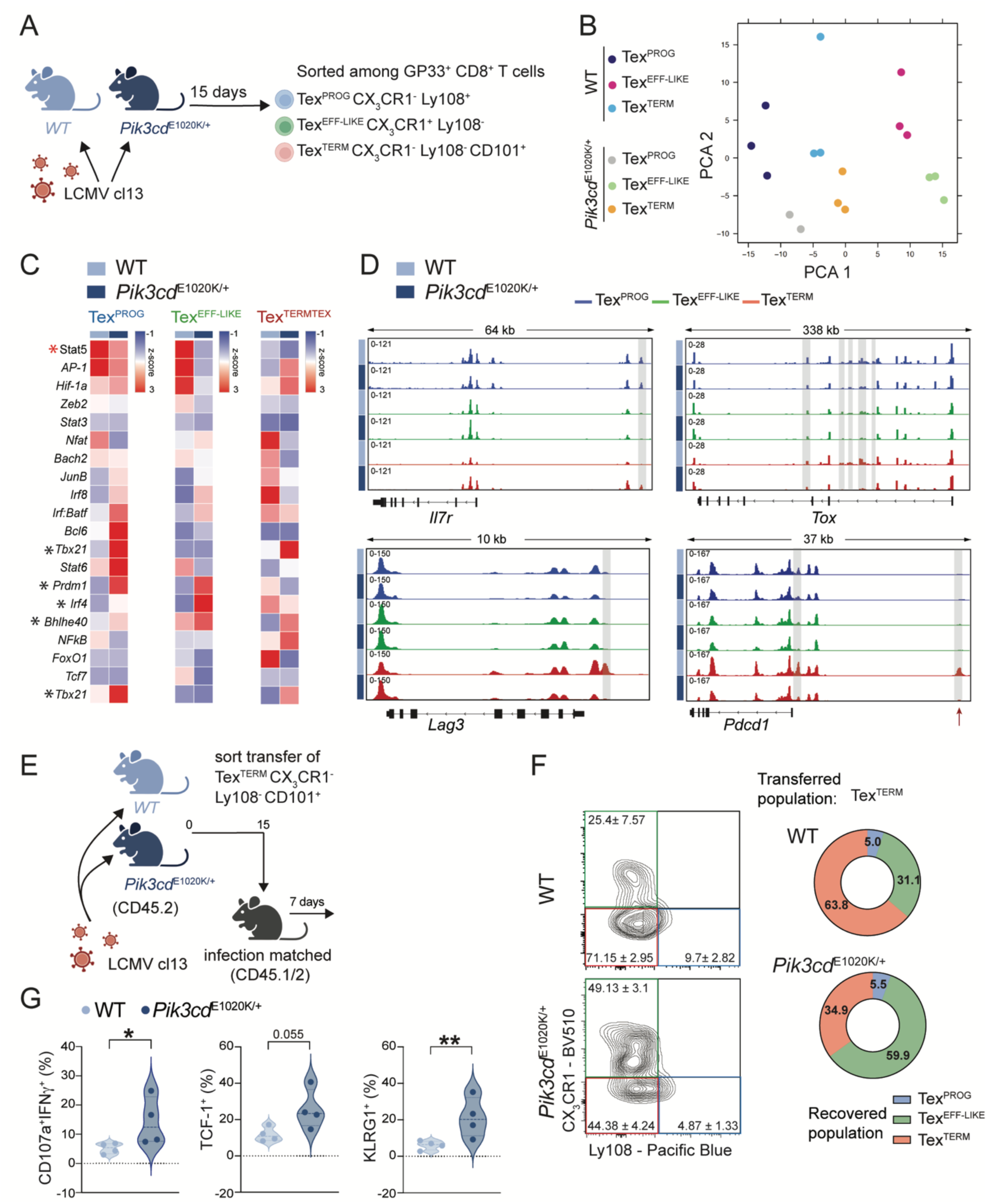
Activated PI3Kδ prevents epigenetic changes associated with exhaustion and promotes plasticity. **(A-D)** GP33^+^ CD8^+^ antigen-specific Tex^PROG^, Tex^EFF-LIKE^ and Tex^TERM^ cells were sorted from C57BL/6 WT and *Pik3cd*^E1020K/+^ mice d15 p.i. with LCMV cl13 and analyzed by ATAC-seq. **(A)** Experimental design**. (B)** Principal component analyses (PCA) of indicated subpopulations. **(C)** TF motif enrichment in ChaRs compared between WT and *Pik3cd*^E1020K/+^ among Tex^PROG^, Tex^EFF-LIKE^ and Tex^TERM^ cells. Black Asterix indicate motifs of interest, red star indicates STAT5 motif enriched in all 3 WT Tex subpopulations. **(D)** ATAC-seq tracks at selected genes (*Il7r*, *Tox, Lag3* and *Pdcd1*) from Tex^PROG^, Tex^EFF-LIKE^ and Tex^TERM^ (top to bottom). Shaded boxes indicate peaks with significantly different accessibility between WT and *Pik3cd*^E1020K/+^. The red arrow indicates the *Pdcd1* ∼23-kb enhancer. **(E-G)** d15 p.i. sorted Tex^TERM^ cells (50-100x10^3^) from C57BL/6 WT or *Pik3cd*^E1020K/+^ mice were transferred into infection matched host. Transferred cells were analyzed d7 post transfer. **(E)** Experimental design. **(F)** Representative FACS plots (left) and donut plots (right) showing percentages of indicated recovered population. Numbers in contour plots indicate means ± SEM of shown experiment. Donut plot showing means of 3 pooled experiments. **(G)** Graphs showing percentages of IFNγ^+^CD107α^+^ post *ex vivo* peptide stimulation (left), TCF-1^+^ (middle) and KLRG1^+^ (right) on recovered transferred cells. For ATAC-seq analyses, cells were sorted from 3 WT and 3 *Pik3cd*^E1020K/+^. Numbers in contour plots indicate mean +/- SEM of shown experiment. In **(G)** each dot represents 1 mouse. Statistical differences were determined Student’s t-test **(G)** *P < 0.05, **P < 0.01, ***P < 0.001.

Analyses of differential ChARs on specific genes provided additional mechanistic insight: for example, a ChAR upstream of *Il7r* was more accessible in activated PI3Kδ Tex^TERM^ cells, similar to markings seen in Tex^PROG^ cells (**Fig.7D**). In contrast, the accessibility pattern of *Tox* was decreased in Tex^TERM^ from *Pik3cd*^E1020K/+^ mice (**Fig.7D**). Although not all genes encoding inhibitory receptors showed altered chromatin accessibility (see **Extended Data Fig.7A** for *Havcr2* and *Cd101*), accessibility of ChARs associated with *Lag3* and *Pdcd1*, including a well-characterized exhaustion-specific enhancer in *Pdcd1* ∼23 kilobases (kb) upstream from the transcriptional start site, were significantly decreased in Tex^TERM^ from activated PI3Kδ mice compared to WT (**Fig.7D**). Deletion of this enhancer leads to increased effector-like cell generation ^59^.

The decreased expression of TOX and lack of epigenetic changes associated with exhaustion suggested that *Pik3cd*^E1020K/+^ cells may not be fully committed to a terminally-exhausted state. To evaluate plasticity of Tex^TERM^ cells, Ly108^-^CX_3_CR1^-^CD101^+^ Tex^TERM^ cells were sorted from d15 LCMV cl13 infected mice, and transferred into infection-matched WT recipients that were analyzed 7 days later **(Fig.7E and Extended Data Fig.7B-C)**. The bulk of Tex^TERM^ cells (63.8 **±** 5.8%) remained Tex^TERM^ cells when transferred from WT mice (**Fig.7F**). However, a smaller percentage (34.9 **±** 7.5%) of Tex^TERM^ cells from activated PI3Kδ mice remained as Tex^TERM^ cells, whereas approximately 60 **±** 6.5 % differentiated into CX_3_CR1^+^ Tex^EFF-LIKE^ cells, acquiring expression of CX_3_CR1 and KLRG1 as well as increased expression of effector cytokines (**Fig.7F-G)**. In line with increased progenitor signature in Tex^TERM^ from *Pik3cd*^E1020K/+^ mice, these cells also displayed higher TCF-1 expression post transfer compared to WT cells **(Fig.7G)**. Together, these results establish a role for PI3Kδ activation in both promoting the differentiation of effector cells and preventing the epigenetic fixing of Tex^TERM^ cells, thereby allowing both functional plasticity and persistence during chronic infection.

## DISCUSSION

Understanding the pathways that shape T cell responses during persistent antigen exposure is an important goal for reinvigorating T cells during chronic infection and cancer. Although several lines of evidence have pointed to roles for PI3Kδ and its effectors in regulating factors important for T cell exhaustion ^21, 22, 23, 25, 26, 31, 35, 60, 61^, the direct consequences of PI3Kδ activation in the context of chronic infection have not been examined *in vivo*. Using transcriptomic, epigenomic, functional assays and a mouse model expressing activated PI3Kδ, we provide evidence that PI3Kδ is a critical hub that integrates multiple inputs to regulate cell fate during chronic infection. While activated PI3Kδ was deleterious for the maintenance of TCF-1^+^ progenitor cells, PI3Kδ activation promoted effector cell differentiation and function during chronic infection, suppressing TOX expression, preventing terminal epigenetic imprinting and allowing T cells with effector function to be maintained, with improved viral clearance over time.

Recent studies show that expression of FoxO1 in CAR-T cells provides a selective advantage in part by promoting TCF-1 expression without attenuating effector differentiation ^21, 26^. Nonetheless, expression of an AKT-resistant mutant of FoxO1 was seen to “fix” cells in a progenitor state and diminished CAR-T antitumor function, highlighting the requirement for context-dependent regulation by PI3K. Our finding of FoxO1-dependent reduction of Tex^PROG^ cells in *Pik3cd*^E1020K/+^ mice during chronic infection aligns with recent studies showing that inhibition of PI3Kδ increases TCF-1^+^ expression in culture ^28, 60, 61, 62^, PI3K-dependent suppression of FoxO1 ^43^, and studies of FoxO1-deficient CD8^+^ T cells ^23, 25^. However, we also found that activated PI3Kδ inhibited expression of *Myb* and *TOX*, which are critical for Tex^PROG^ maintenance ^8, 9, 10, 11, 45^. Despite these differences, the Tex^PROG^ cells that remained were sufficient and functional to maintain LCMV-specific responses and viral control, which was associated with expanded effector-like cell populations, reminiscent of the increased EBV-specific cells seen in APDS1 ^63^. While TCF-1-deficiency in CD8^+^ T cells also promotes effector cell phenotypes ^4^, loss of TCF-1 prevents the maintenance of antigen-specific T cells during chronic infection ^3^. These findings suggest a more complex picture of the regulation of T cell persistence in the presence of chronic antigen exposure.

Although FoxO1 is a major TF inhibited by PI3Kδ, effector cell expansion was largely independent of FoxO1-inactivation, at least at early time points of chronic infection. Indeed, multiple PI3K effectors influence exhaustion, including TFs such as Bach2, which plays a key role in promoting Tex^PROG^ cells ^24^ and KLF2, which suppresses features of exhaustion ^22^, as well as regulators of mTORC1 ^64^, and FoxO1- and AKT-independent effectors. Together, these factors may contribute to the broad effects of activated PI3Kδ.

Mechanistically, *Pik3cd*^E1020K/+^ mice had more *Il21*-expressing T_FH_ cells 7d p.i., highlighting key roles for T_FH_ in the regulation of multiple cell populations during infection ^40, 65^, as well as increased sensitivity to IL-21, with elevated STAT3 phosphorylation (pSTAT3) and survival of *Pik3cd*^E1020K/*+*^ CD8^+^ T cells. In CD8^+^ T cells, this may result from increased *Il21r* and decreased *Socs1* and *Socs3*; however, increased pSTAT3 is also seen in fibroblasts transformed by PI3Kα, suggesting broad effects of PI3K on this pathway ^66, 67^. Moreover, our epigenetic analyses revealing increased BATF:IRF and BLIMP1 motif-binding in *Pik3cd*^E1020K/+^ CD8^+^ T cells, tie closely with data showing IL-21 induces BATF and BATF:IRF4 complexes that support BLIMP1, (which itself is induced by STAT3 ^58^), to promote effector cell function ^68^. These observations, along with the increased cell proliferation signatures seen in *Pik3cd*^E1020K/+^ cells on d8 p.i. and effects of IL-21 on CD8^+^ T cell survival, may account for the persistence of CD8^+^ T cells and increased viral clearance, two key goals of immunotherapy, despite the reduced expression of TCF-1. It is of note that PD-1 recruits phosphatases SHP-1 and SHP-2 which antagonize key signaling molecules downstream of the TCR and CD28 co-stimulation, leading to inhibition of PI3K/AKT cascades ^69, 70^. Thus, PD-1 immune checkpoint blockade enhances effector function in CD8^+^ T cells in part by increasing PI3K signaling, and mice expressing kinase-inactive PI3Kδ fail to respond to anti-PD-1 blockade ^33^, although low dose PI3Kδ inhibition has also been reported to potentiate ICB responses, likely by promoting TCF-1^+^ progenitor cells ^62, 71^. It is therefore of interest that two recent base-editing screens designed to detect mutations that promote CD8^+^ and CAR-T cell function uncovered multiple activating mutations affecting PI3Kδ as top hits ^34, 35^. Together, these observations highlight the importance of PI3K pathways for promoting T cell activation during exhaustion.

Both IL-2/STAT5 and STAT3 pathways have been implicated in driving effector cells in chronic infection ^72, 73^. However, despite our previous findings of increased STAT5 signatures in cultured *Pik3cd*^E1020K/+^ CD8^+^ T cells ^27^, IL-2/STAT5 chromatin markings were strongly enriched in WT cells. In contrast, STAT3 signatures were prominent in *Pik3cd*^E1020K/+^ CD8^+^ T cells, likely due to the increase in IL-21-producing T_FH_ cells. It is also of interest that we observed increased IFNα signatures in WT relative to *Pik3cd*^E1020K/+^ Tex cells. This observation parallels findings in CD4^+^ T cells, where Type I IFNs have been reported to be associated with elevated IL-2/STAT5 responses, which counteract STAT3 signals that drive T_FH_ cell differentiation ^74^. These observations suggest that activated PI3Kδ influences differentiation decisions, in part, by affecting both production of and responses to different cytokines, resulting in distinct effector T cell populations.

Although Tex cells are thought to be terminally committed to their dysfunctional exhausted fate, a large fraction of Tex^TERM^ from *Pik3cd*^E1020K/+^ mice produced cytokines and acquired CX_3_CR1^+^effector cell characteristics in cell-transfer experiments. While Tex^TERM^ cells in *Pik3cd*^E1020K/+^ mice cluster with those from WT mice, the repression of TOX and lack of characteristic chromatin markings at loci associated with “epigenetic scarring” in *Pik3cd*^E1020K/+^ Tex^TERM^ cells suggests that PI3K activation protects against or prevents a fixed state of terminal exhaustion. Although lower viral titers or altered TCR usage may contribute to these phenotypes, P14 co-transfer experiments provided evidence that these phenotypes were seen even with a fixed TCR in the same environment as WT cells; similar phenotypes were also seen early in infection, prior to the reduction in viral titers. Indeed, both Tex^TERM^ and Tex^PROG^ from *Pik3cd*^E1020K/+^ mice preferentially differentiate into Tex^EFF-LIKE^ cells, perhaps reflecting a more plastic state induced by elevated PI3Kδ activity. Alternatively, heterogeneity in Tex^TERM^ cells may be more pronounced in the presence of activated PI3Kδ and indeed both Tex^TERM^ and Tex^PROG^ showed increased effector gene signatures.

While multiple studies have underscored the role of key TFs in regulating population dynamics during exhaustion, our work highlights the central role of PI3Kδ signaling as a master regulator of effector function and persistence, with potential therapeutic implications. In particular, while PI3Kδ inhibitors may be useful to maintain a robust TCF-1^+^ population in cultured CAR-T or other T cell populations *in vitro* ^28, 61^, activation of PI3Kδ via cytokines or other approaches *in vivo* may be critical for CD8^+^ T cells to develop into effector populations, via both cell-intrinsic and extrinsic mechanisms. It is therefore of interest that although PI3Kδ inhibitors have had mixed success treating tumors, recent studies have shown promising results using intermittent PI3Kδ inhibition in mice ^75, 76^. While much of this work has focused on effects on Tex^PROG^ and T_REGS_, our studies suggest this type of approach may permit full activation of CD8^+^ effector T cell potential. Our results suggest that PI3Kδ is a dual-edged sword and that careful manipulation of these pathways may allow for more refined approaches for reinvigorating T cells during exhaustion.

## METHODS

### Mice

Mice were maintained and treated under specific pathogen free (SPF) conditions in accordance with the guidelines of NHGRI (protocol G98-3), NIAID (protocol LISB-22E), NINDS (protocol 1295-23) and the Animal Care and Use committees at the NIH (Animal Welfare Assurance #A-4149-01). *Pik3cd*^E1020K/+^ mice have been described previously and were backcrossed > 10 times to C57BL6/J ^27, 36^. Age and sex matched mice between 8-12 weeks of age were used. For adoptive transfer experiments, congenic CD45.1/2 (C57BL/6J x B6.SJL-*CD45a(Ly5a)/Nai F1)* mice were obtained from NIAID-Taconic contract facility and used as recipients. C57BL/6J expressing the P14 TCR transgene (JAX: Tg (TcrLCMV)327Sdz) were crossed to *Pik3cd*^E1020K/+^ to generate donor mice. OT-1 *Pik3cd*^E1020K/+^ mice have been previously described ^27^.

### Infections

LCMV clone 13 and Armstrong viral stocks were maintained by Dorian McGavern. Mice were infected with 2x10^5^ plaque-forming units (PFU) of LCMV Armstrong strain intraperitoneal (ip) or intravenously (iv) with 2x10^6^ PFU of LCMV clone 13 by tail vein injection ^3^. Viral titers in the serum, lung, liver and kidney were determined by plaque assay using Vero cells ^3^. For survival analysis, mice were monitored daily according to institutional ethic guidelines and were euthanized when mice developed signs of reduced mobility, hunched posture, or respiratory distress. Clinical score was assessed as previously described ^77^.

### In vivo treatments

For CD4 depletion, mice were injected with 500µg anti-CD4 depleting mAbs (clone GK1.5, BioXcell) 1 day before and 1 and 3 days post-infection. For IL-21R blockade, mice were injected with 500µg anti-IL21R blocking mAbs (clone 4A9, BioXcell) 1 day before and 3 days after infection. Control groups received 200µg of rat IgG2b (clone LTF-2, BioXcell).

### Tissue processing and lymphocyte isolation

Livers and lungs were perfused with PBS. Peripheral blood mononuclear cells were isolated from spleen, liver and lung. Spleens and livers were mechanically homogenized against a 70μM cell strainer using the plunger of a 3ml syringe. Lungs were cut into pieces, incubated with digestion cocktail containing 0.25mg/ml liberase in HBSS buffer for 45 min at 37°C. Cell suspensions were filtered onto a 70μM cell strainer. Cell suspensions from lung, liver and spleen were resuspended in 1-2ml of ACK lysis buffer (GIBCO) for 5 min at room temperature (RT). Lung and liver cell suspension were enriched for lymphocytes by centrifugation over Percoll density centrifugation (37% (v/v) Percoll in PBS) for 30 min at 4°C (500g).

### Adoptive cell transfer

For P14 adoptive transfer experiments, P14 CD8^+^ T cells were isolated from spleen of *Pik3cd*^E2010K/+^ or WT P14 mice using Miltenyi mouse negative selection CD8^+^ T isolation kits. 5x10^3^ isolated P14 cells were injected i.v. into congenic CD45.1/2 or *Pik3cd*^E2010K/+^ recipient mice that were subsequently infected with LCMV cl13. After 8 days, CD8^+^ P14 cells from the spleen were analyzed. Alternatively, *Pik3cd*^E1020K^or WT littermates were infected with LCMV cl13. At d15, mice were euthanized, spleens collected and enriched for CD8^+^ T cells using negative selection Miltenyi mouse CD8^+^ T isolation kits. Cell suspensions were stained as described previously and exhausted CX_3_CR1^-^Ly108^+^CD101^+^ Tex^TERM^ and Ly108^+^CX_3_CR1^-^ Tex^PROG^ among live CD3^+^CD8^+^PD-1^+^CD44^+^ T cells were sorted with FACSaria^TM^ Fusion (BD biosciences). 50-100 x 10^3^ cells were transferred into infection match CD45.1/2 recipient mice. At d8 post-transfer, spleen and liver were collected and CD45.2 CD8^+^ T cells analyzed.

### Retroviral transduction

For retrovirus production, HEK 293T cells were co-transfected with GFP-pMIGR or Foxo1-WT-pMIGR (2mg) along with pCL-Eco helper plasmid (1mg). 48 hrs post-transfection, cell supernatants were collected and incubated with 5x PEG-IT reagent (SBI) overnight at 4°C. Precipitated retrovirus was centrifuged at 1500xg for 30 min and supernatants were discarded. Retrovirus was centrifugated again at 1500xg for 5 min, supernatant discarded and concentrated retrovirus was reconstituted in complete IMDM. For retroviral transduction, naïve P14 cells were activated with 0.02µg GP33 peptide/ml in a 48-well plate in 1ml of complete RPMI media (10% FBS, 5% Penicillin/Streptomycin, 5% Glutamine and 1µM 2-mercaptoethanol). 18 hrs post-activation concentrated retrovirus (20μL) was mixed with polybrene A (8µg/ml final), added to 1ml of activated P14 cells in a 48 well plate and spinfected at 2000rpm for 1 hr at 37°C). Transduced cells were subsequently cultured at 37°C for a total of 72 hrs. Transduced cells were detected by flow cytometry using GFP expression.

### CD8^+^ T cell gene editing

Ribonucleoprotein (RNP) complexes were assembled using recombinant Cas9 (IDT) and predesigned Alt-R CRISPR-Cas9 gRNAs (IDT) duplexed with atto550 conjugated tracrRNA (IDT). CD8^+^ T cells were isolated from the spleen using mouse CD8^+^ T cell isolation kit (Miltenyi Biotec). 10x10^6^ CD8^+^ T cells from P14 mice were resuspended in 20µL P3 primary cell nucleofection buffer (Lonza), mixed with 5uL of Cas9-RNP complex and transferred to wells of a 16 well nucleocuvette strip (Lonza-V4XP-3032). P14 CD8^+^ T cells underwent nucleofection using the DN100 program in a Lonza 4D-nucleofector device. Immediately after nucleofection, cells were harvested and cultured in complete RPMI supplemented with IL-7 (10 ng/mL) for 24 hrs. Following incubation, cells were harvested and washed. 5x10^3^ P14 CD8^+^ T cells were adoptively transferred into recipient mice, which were infected with LCMV cl13 and analyzed at d7 post infection. For negative control treated cells, RNP complexes included negative control crRNA #1 (IDT). For *foxo1* targeted cells, RNP complexes included three predesigned sgRNAs (IDT) targeting the *foxo1* gene. Guides: Mm.Cas9.FOXO.AA – CCACTCGTAGATCTGCGACA; Mm.Cas9.FOXO.AB – CGGGTCGGTCTCCACCACCT; Mm.Cas9.FOXO.AC – CAACTCGACCACCTCCAGTC; Negative control scrRNA #1 – cat# 1072544 (IDT) – no guide sequence publicly available.

### Antibodies and Flow cytometry

Single cells suspensions were obtained from spleen, lung and liver. Before surface staining, cells were incubated with fixable viability dye (Thermo Fisher) for 10 min at 4°C. Single cell suspensions were then stained for surface markers for 45-60 min at 4°C in flow cytometry buffer (PBS, 2%FBS, 2 mM EDTA) with the following antibodies: CD8 BUV395 (53-6.7), CD4 BUV496 (GK1.5), CD45.1 PE (A20), CD45.1 BUV737 (A20), Ly108 BV650 (13G3), Tim3 PE-CF594 (5D12) (BD Biosciences), CD3 FITC (17A2), CD3 APC/Fire^TM^810 (17A2), CD45.2 Alexa Fluor® 700 (104), CD107a (LAMP1) APC (1D4B), Ly108 Pacific Blue^TM^ (330-AJ), PD-1 BV605 (29F.1A12), PD-1 BV737 (29F.1A12), CD62L BV711 (MEL-14), CD223 (LAG3) BV785 (C9B7W), CX_3_CR1 BV510 (SA011F11) (Biolegend), CD4 PE (GK1.5), CD39 SuperBright^TM^436 (24DMS1), CD44 AlexaFluor^TM^ 700 (IM7), CD223 (LAG3) PerCP-eFluor^TM^710 (3DS223H), CD101 PE-Cyanine7 (Moushi101), KLRG1 PE-Cyanine7 (2F1) (Thermo Fisher) (see **Supplementary Table 9).** LCMV-derived D^b^ GP_33-41_ (KAVYNFATM) tetramer-APC and LCMV-derived I-A^b^ GP_66-77_ (DIYKGVYQFKSV) tetramer-PE were obtained from NIH tetramer core facility. GP33 and GP66 tetramer was stained together with cell surface markers. For intracellular staining for transcription factors, cells were fixed and permeabilized using BD Cytofix/Cytoperm permeabilization kit (BD Biosciences) or Transcription Factor Fixation/Permeabilization kit (Thermofisher) with the following antibodies: FoxO1 Pacific Blue (C29H4), Tox 488 (E6G50), TCF-1 AF647 (C63D9), Ki67 AF532 (SolA15), granzyme B APC/Fire^TM^750 (QA16A02), Eomes PerCP-eFluor^TM^710 (Dan11mag), Foxp3 AF700 (FJK-16s), STAT3 PE (4G4B45) (see **Supplementary Table 9**).

Staining of phospho-proteins was performed using fixation in 4% PFA (25 min, RT) followed by permeabilization using ice-cold 100% methanol (1 hr, 4°C) and intracellular staining in 0.5% Triton X-100, 0.1% BSA in PBS (at least 1 hr, 4°C): phospho-STAT3 (Tyr705) (LUVNKLA), PE-Cyanine7 (Thermo Fisher) and phospho-FoxO1 unconjugated (S256) (see **Supplementary Table 9**).

Unconjugated antibodies were detected with secondary antibodies conjugated with AF488. Intracellular cytokine staining was performed after re-stimulation with GP_33-41_ LCMV peptide (AnaSpec) (0.5µg/ml) or anti-CD3 (1µg/ml, BioXCell) and anti-CD28 (5µg/ml, BioXCell) in the presence of GolgiStop (monensin) and CD107a APC (Biolegend) for 90-150 minutes at 37°C. Intracellular staining for IFNγ BV605 (XMG1.2) or IFNγ BV786 (XMG1.2) and TNFα BV421 (MP6-XT22) (BD Biosciences) was performed with FoxP3/transcription factor staining buffer set (ebioscience). Data were collected with Fortessa X20 flow cytometers (BD Biosciences) or Cytec Aurora (CytekBioscience) and analyzed with FlowJo software (TreeStar).

### Cell culture

OT-1 (2.5x10^6^) splenocytes were stimulated with 10nM OVA_257-264_ (AnaSpec) for 3 days in complete media ^27^. Cells were rested in serum-free media for 1hr before IL-21 was added to culture (Thermofisher, 40ng/ml). Alternatively, CD8^+^ T cells were activated for 3 days in complete media with anti-CD3 (1μg/mL) and anti-CD28 (3μg/mL) for 3 days with or without IL-21 (Thermofisher, 40ng/ml).

### Measurement of liver cytokines and chemokines in tissue homogenates

Left liver lobes were isolated and added to an ice-cold mixture of 500μL RIPA buffer supplemented with protease inhibitor. Using Bertin hard tissue grinding tubes, samples were homogenized for 2 cycles of 20 seconds at 5500rpm in a Precellys Evolution (Bertin). Homogenates were centrifuged at 8000xg for 10 min to remove debris and supernatants were collected. Cytokine and chemokines were measured in homogenates using a 32plex Luminex kit (Millipore) and a Bio-Plex 200 analyzer (BioRad).

### Bulk RNA sequencing

*Pik3cd*^E2010K/+^ or C57BL/6 WT littermates (n=3 for each group) were infected with LCMV cl13 (iv, 2.10^6^ pfu). Antigen-specific CD8^+^ T cells were sorted directly into Trizol (1mL final volume) at d8 p.i.. Total RNA was isolated from samples using phenol-chloroform extraction with GlycoBlue as a co-precipitant. RNA was quantified using Qubit RNA high sensitivity reagents and 30ng of total RNA was used as input for RNAseq library preparation. The quality of the isolated total RNA from each sample was checked on 2200 Tapestation System (Agilent, Santa Clara, USA) using the RNA ScreenTape assay and the quantitation was performed using the NanoVue Plus Spectrophotometer (Biochrom Ltd). 100-300ng of total RNA per sample with a RNA integrity number >8 were used for library preparation. The RNA-Seq library preparation was performed using NEBNext poly(A)+ mRNA magnetic isolation module and NEBNext Ultra RNA Library prep kit. Libraries were sequenced using the NovaSeq platform (Illumina) and underwent paired-end sequencing to produce between 262 and 540 million 100bp readpairs per sample, for a total of 13.8 billion read pairs. Adapter sequences and low-quality bases were trimmed using Cutadapt v1.18, followed by alignment with the reference genome (mm10) and annotated transcript using STAR v2.7.0f. The gene expression quantification analysis was performed using RSEM v.1.3.1. Differential gene expression was performed in R 4.2.1 using DESeq2 v 1.36.0. Genes with low expression were pre-filtered (genes with at least 10 counts in two or more samples were retained). Absolute fold change > 1.5 and adjusted p-value < 0.1 were classified significant. Volcano plots were created using ggplot2 v3.3.6, to visualize the differentially expressed genes.

### Single CD8 cell preparation and RNA sequencing

Antigen-specific CD8^+^ T cells were sorted from spleens of infected *Pik3cd*^E2010K/+^ and C57BL/6 WT littermates on d8 and d28 LCMV cl13 p.i. (n=2-6 for each group per experiment). Single cell suspensions were prepared as previously described. Live antigen-specific CD8^+^ T cells were sorted using Aria Fusion cell sorter (BD). Labeled cells were pooled by genotype for d28 p.i. sequencing (2 replicates per genotype) or labeled with unique Hashtag antibodies (Biolegend) and pooled together for d8 p.i. sequencing. CD8^+^ T cell from mice d28 p.i., cells were pooled together by genotype with 1x10^4^ cells per donor used as input. Chromium Single Cell 3’ v2 libraries (10x Genomics) were generated following the standard manufacturers protocol and sequenced on one NovaSeq XPlus (d8) or NovaSeq 600 (d28) run at the Frederick National Laboratory for Cancer Research Sequencing Facility. The sequencing run was setup as a 28 cycles + 91 cycles non-symmetric run. All samples had sequencing yields of more than 206 million reads per sample. For d28 p.i., the *Pik3cd*^E2010K/+^ samples had 853 and 921 million reads while the WT samples had 865 and 687 million reads. The Q30 values of *Pik3cd*^E2010K/+^ samples were approximately 97%, 93%, and 96% for the barcode, RNA read, and UMI, respectively. The Q30 values of WT samples were approximately 97%, 93%, and 96% for the barcode, RNA read, and UMI, respectively. Median genes per cell were 2,001 and 2,088 for the *Pik3cd*^E2010K/+^ samples and 2,339 and 2,436 for the wildtype samples. Sequencing saturation for all the samples were between 90% and 93%. For the TCR sequencing data on these samples, the cellranger (version 7.1.0) vdj command was used with vdj_GRCm38_alts_ensembl-7.0.0 reference. For d8 p.i, the three multiplexed libraries had 2.5, 2.9 and 3.9 billion reads. The Q30 values for the libraries were above 96.2%, 94.7%, and 97.4% for the barcode, RNA read, and UMI, respectively. Median genes per cell were 3,883, 4,117, and 3,967 for the three replicates. Sequencing saturation were 87%, 88.3% and 89.9%. The raw data were processed using cellranger 7.1.0 (d28) or cellranger 7.1.0 (d8) which included alignment to the mouse reference mm10-2020-A with the include introns parameter set to True. The samples were integrated using Harmony which was performed using IntegrateLayers function with method set to HarmonyIntegration.

### Single CD4 cell preparation and RNA sequencing

For total CD4^+^ single cell sequencing, spleens from cl13 infected 4 *Pik3cd*^E2010K/+^ or C57BL/6 2 WT littermates were collected at d8 p.i. and single cell suspension were prepared as described above. Live total CD4^+^ T cells were sorted using Aria Fusion cell sorter (BD). CD4^+^ T cells from d8 p.i. were labeled with unique Hashtag antibodies (Biolegend). Labeled cells were pooled by genotype. For CD4 d8 sequencing, multiplexed libraries of sorted total CD4^+^ T cells from 4 *Pik3cd*^E2010K/+^ and 2 WT were pooled by genotype and processed using hashtag barcoding, with 1x10^4^ cells per donor used as input. Chromium Single Cell 5’ v2 libraries (10x Genomics) were generated following the standard manufacturers protocol and sequenced on one NovaSeq XPlus (d8) run at the Frederick National Laboratory for Cancer Research Sequencing Facility. The sequencing run was setup as a 28 cycles + 91 cycles non-symmetric run. All samples had sequencing yields of more than 206 million reads per sample. The Q30 values of WT and *Pik3cd*^E2010K/+^ samples were approximately 97%, 93%, and 96% for the barcode, RNA read, and UMI, respectively. Initial processing was accomplished using cellranger 7.0.1 which included alignment to the mouse reference mm10-2020-A with the include introns parameter set to True. Median genes per cell were 1662 and 1854 for the *Pik3cd*^E2010K/+^ samples and 1917 and 1759 for the WT samples. Sequencing saturation for all the samples were between 87% and 88%.

### Single cell RNA-seq bioinformatic analysis

For CD4^+^ data, In the downstream analysis of the expression data, the Cell Ranger aggregation output (filtered_feature_bc_matrix directory) was loaded into Seurat version 5.1.0^78^ using the CreateSeuratObject function with min.cells=3. For filtering out low-quality cells, we established minimum thresholds for gene (nCount_RNA > 1000) and UMI counts (nFeature_RNA > 300), and a maximum threshold for percentage mitochondrial content (percent.mt < 7) based on visual inspection of the corresponding metric distributions across all cells. Log-normalization (LogNormalize function) of the RNA count data and centered log-ratio transformation (CLR) of the HTO count data were carried out using the NormalizeData function. For HTO demultiplexing cells into the individual mice and removing any doublets, HTODemux was run with a positive quantile = 0.99. The top 2000 variable features were determined with the VariableFeatures function, expression data was scaled with ScaleData, and dimensionality reduction was performed with the RunPCA function on the variable features, followed by determining the dimensionality of the dataset on an elbow plot (ElbowPlot function), which was determined to be 40. Subsequently, FindNeighbors, FindClusters, and RunUMAP were run on 40 dimensions and a range of resolutions to determine the optimal resolution that captures biological diversity in the dataset, which was determined to be 0.5, resulting in 10 clusters. The clusters identified by the FindClusters function of Seurat (default Louvain clustering setting and 0.25 resolution) were annotated using cell type-specific markers. This clustering approach yielded two sets of mirrored clusters with similar cell type compositions. Upon comparison of both sets of clusters, we identified one of the mirrored set of clusters as low quality cells due to their higher percentage mitochondrial genes, high percentage of ribosomal genes, and lower percentage of total counts. Further analysis revealed that cluster with low quality cells produced similar outcomes in terms of results when compared to clusters with higher quality. Accordingly, further analysis was performed only on the set of clusters with higher quality metrics, which were reclustered as described above (starting at the FindVariableFeatures step) with 45 dimensions and a resolution of 0.75, resulting in 9 clusters, which were annotated using cell type-specific markers.

For CD8 d8 downstream analysis following Cell Ranger processing was performed in Seurat 5.2.0. The cells were demultiplexed using Seurat HTODemux function followed by normalizing the HTO assay using CLR method. The singlet cells were filtered using median absolute deviation of number of the features and mitochondrial reads lower than 5%. After filtering, data were log-normalized using NormalizeData, top 2000 variable features were determined using FindVariableFeatures, scaled using ScaleData, and obtained principal component using RunPCA. The samples were integrated using Harmony which was performed using IntegrateLayers function with method set to HarmonyIntegration. Nearest neighbor graph was constructed using 30 dimensions of harmony. Cells were clustered using a graph-based approach via the FindClusters function in Seurat with resolution parameter set to 0.2. FindAllMarkers function was used to find marker genes for the clusters. The cells were identified based on the cluster markers. DE genes were identified using FindMarkers on each celltype using Seurat default parameters.

For CD8 d28, the downstream analysis is performed in Seurat 4.3.0. Cells with number of features lower than 500 and higher than 4000 and mitochondrial reads higher than 5% were filtered out. Cells were annotated by SingleR 2.4.0 using celldex ImmGenData reference data. Non-T cells were filtered out from downstream analysis. After filtering, data were log-normalized using NormalizeData. The samples were CCA-integrated using the FindIntegrationAnchors and IntegrateData functions. The integrated assay was scaled using ScaleData and obtained principal component using RunPCA. Based on elbow plot, 30 principal components were used for FindNeighbors and subsequently FindCluster functions. Cluster markers and DE genes were identified using FindAllMarkers and FindMarkers functions, respectively with default parameters. To examine the exhaustion module score of progenitor, effector-life and terminal exhausted T cells in all clusters (**Supplementary Table 2**), we used the AUcell R package ^79^. Briefly, we converted our data into an expression matrix of read counts per cell and identified whether genes contained in the exhaustion module score gene set were expressed in each cell. We then used AUcell to generate an enrichment score of Tex^PROG^, Tex^EFF-LIKE^ and Tex^TERM^ in all individual cells compared to all other genes. To visualize these results, we used Seurat to plot enrichment p-values as a feature plot.

Trajectory analysis was performed using slingshot 2.10.0 setting 𝑇𝑒𝑥_*PROG*_ as the starting clusters (start.clus). Barplots are created using ggplot2 3.4.4 and heatmaps are created using ComplexHeatmap 2.18.0. TCR repertoire sequencing data were analyzed using scRepertoire 1.10.1. The count data were read from the cellranger output ‘filtered_contig_annotations’ of four libraries and were combined using combineTCR function and were merged with the scRNA-seq Seurat object using combineExpression. Each clonotype was defined by the amino acid sequence of the paired CDR3 regions of the 𝛼 and 𝛽(TRA-TRB) chains. The clonalQuant function was used for quantification of abundance setting scale parameters TRUE. Using ggplot2 v3.4.4, the frequency of top 10 clonotypes of each celltype was plotted.

For CD8^+^ T cell d28 p.i pseudobulk analysis, the raw counts were aggregated across cells for each biological replicate on the indicated clusters using AggregateExpression. Genes with a very low level of expression across biological replicates, less than 10 total counts across all samples, were filtered from the dataset. Differential gene expression analysis was performed using DESeq2 v1.44.0^80^, using default parameters, and not including a log2 fold change shrinkage step. The normalized counts were extracted from the DESeq2 object and used for visualization along the corresponding adjusted p-values and log2 fold changes. Gene list can be found in **Supplementary Table 6**. Seurat v4.1.1 was used to generate dimension plots showing clusters and cell types of these data, as well as to generate features plots of gene expression levels. Enriched functional gene sets were identified using ShinyGo^81^ and Enricher^82^, with differentially expressed genes for indicated comparisons used as input.

### ATACseq

*Pik3cd*^E2010K/+^ or WT littermates were infected with LCMV cl13. At the d15 p.i., antigen-specific CD8^+^ T cells were isolated from the spleen using mouse CD8^+^ T cell negative isolation kit (Miltenyi Biotec) and live CD8^+^GP33^+^ Tex^PROG^ (Ly108^+^CX_3_R1^-^), Tex^EFF-LIKE^ (CX_3_CR1^+^Ly108^-^) and Tex^TERM^ (CX_3_CR1^-^Ly108^-^CD101^+^) T cells were sorted with FACSaria™ Fusion (BD biosciences). Briefly, 10 000 cells pooled from 3 independent mice were lysed in ice-cold lysis buffer and the transposition reaction was performed using the Tn5 transposase at 37°C for 30 min. DNA was purified using the QIAGEN MinElute kit (QIAGEN). The libraries were prepared using the Nextera DNA Library Prep Kit (Illumina) and purified using AMPure XP beads (Beckman) following a double-sided protocol to remove primer dimers and large fragments. Samples were performed in triplicates, multiplexed and sequenced on NextSeq-500 (75 bp paired-end reads).

### ATAC seq Bioinformatic analysis

ATACseq reads were processed using our chrom-seek (1.0.2) pipeline (https://github.com/OpenOmics/chrom-seek). In brief, reads were trimmed with Cutadapt version 4.4 (Martin 2011) and then aligned the Mouse mm10 genome using BWA version 0.7.17 ^83^. All reads aligning to the Encode mm10 blacklist regions ^84^ were identified and removed with Picard SamToFastq (https://broadinstitute.github.io/picard/). Reads with a mapQ score less than 6 were removed with SAMtools version 1.17 ^85^ and PCR duplicates were removed with Picard MarkDuplicates. Data was converted into bigwigs for viewing and normalized by reads per genomic content (RPGC) using deepTools version 3.5.1 ^86^.

Peaks were called using Genrich version 0.6 (https://github.com/jsh58/Genrich) and then Differential peaks were called using DiffBind v2 (Stark and Brown, 2011) and its Deseq2 ^80^ differential caller with default parameters. Peaks were then annotated using UROPA version 4.0.2 ^87^ and Gencode Release M18 (GRCm38, Ensembl 93). Motif analysis was performed on differential peaks using HOMER ^57^with default parameters.

## STATISTICAL ANALYSIS

Statistical analyses were performed using GraphPad Prism 9 Software. Mann-Whitney U test or paired and unpaired student t test were used for single comparisons between two groups. For comparison of three or more groups, one-way ANOVA with Tukey’s multiple comparison test, Holm-Sidak multiple test correction, or non-parametric Kruskal-Wallis test with Dunn’s multiple comparison post-test were used. Differences in survival were evaluated with the Mantel-Cox test. P < 0.05 was considered statistically significant (P<0.05 = *; P<0.01 = **; P<0.001 = ***).

### Graphics

Figures were made in Adobe Illustrator 2025 with images generated with GraphPad Prism 10. Drawings were made with Biorender.

## Acknowledgments

We thank V. Lazarevic, R. Bosselut, R. Germain and J.L. Hor for reading and providing helpful suggestions on the manuscript. This research was supported by the Intramural Research Program of the National Institutes of Health (NIH). The contributions of the NIH authors were made as part of their official duties as NIH federal employees, are in compliance with agency policy requirements, and are considered Works of the United States Government. However, the findings and conclusions presented in this paper are those of the authors and do not necessarily reflect the views of the NIH or the U.S. Department of Health and Human Services.

## Author contributions

Study conception and design, A.C.P., J.L.C., and P.L.S.;

Acquisition of data, A.C.P., J.L.C, D.P.G., D.C., J.M.R., S.A., and F.A.O-C.;

Analysis and interpretation of data, A.C.P., J.L.C., S.P, A.D.,P.S., E.A.G., Q.X., and P.L.S.;

Drafting of manuscript, A.C.P., J.L.C. and P.L.S.;

Critical revision and editing, A.C.P., J.L.C., D.C., D.B.M and P.L.S.;

Provision of key materials, D.B.M..

## Data and materials availability

All data, code, and non-commercial materials used in these analyses will be made available upon request. RNA analyses will be available in GEO under submission number GSE299149 and GSE299593.

## Extended Data Figures

**Extended Data Figure 1.**
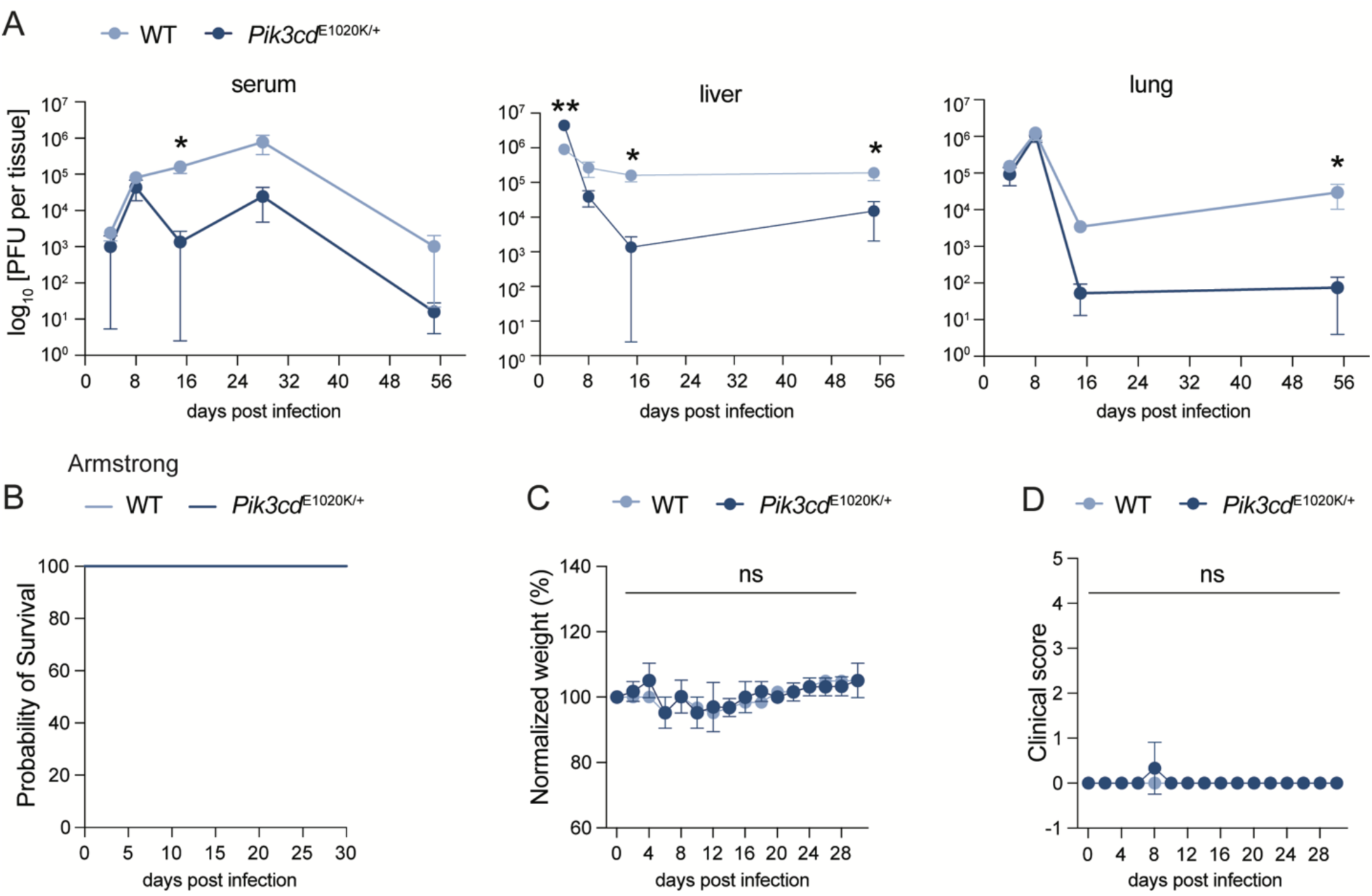
Activated PI3Kδ changes disease outcome during chronic infection. **(A)** C57BL/6 WT and *Pik3cd*^E1020K/+^ mice were infected i.v. with 2x10^6^ PFU LCMV cl13. PFU in serum (left) and liver (middle panel) lung (right panel) at indicated time points. Data are pooled of at least two independent experiments. **(B-D)** C57BL/6 WT or *Pik3cd*^E1020K/+^ mice were infected i.p. with 2x10^5^ PFU LCMV Armstrong. Mice were monitored for survival, weight and clinical score**. (B)** Kaplan Meyer survival curves. **(C-D)** Graphs show normalized weight **(C)** and clinical score **(D)**. Data are representative of two independent experiments. In **(A)** each dot represents pools of 7, 7, 8, and 10 mice on d4, 8, 15, 28 and d55 p.i. in WT mice respectively and 8, 7, 7 and 6 mice on d4, 8, 15, 28 and d55 p.i. in *Pik3cd*^E1020K/+^ mice, respectively. In **(B)** each line represents a pool of 3-4 mice. In **(C)** and **(D)** each dot represents a pool 3 mice. Statistical differences were determined by one-way ANOVA (**A-D**) *p < 0.05.

**Extended Data Figure 2.**
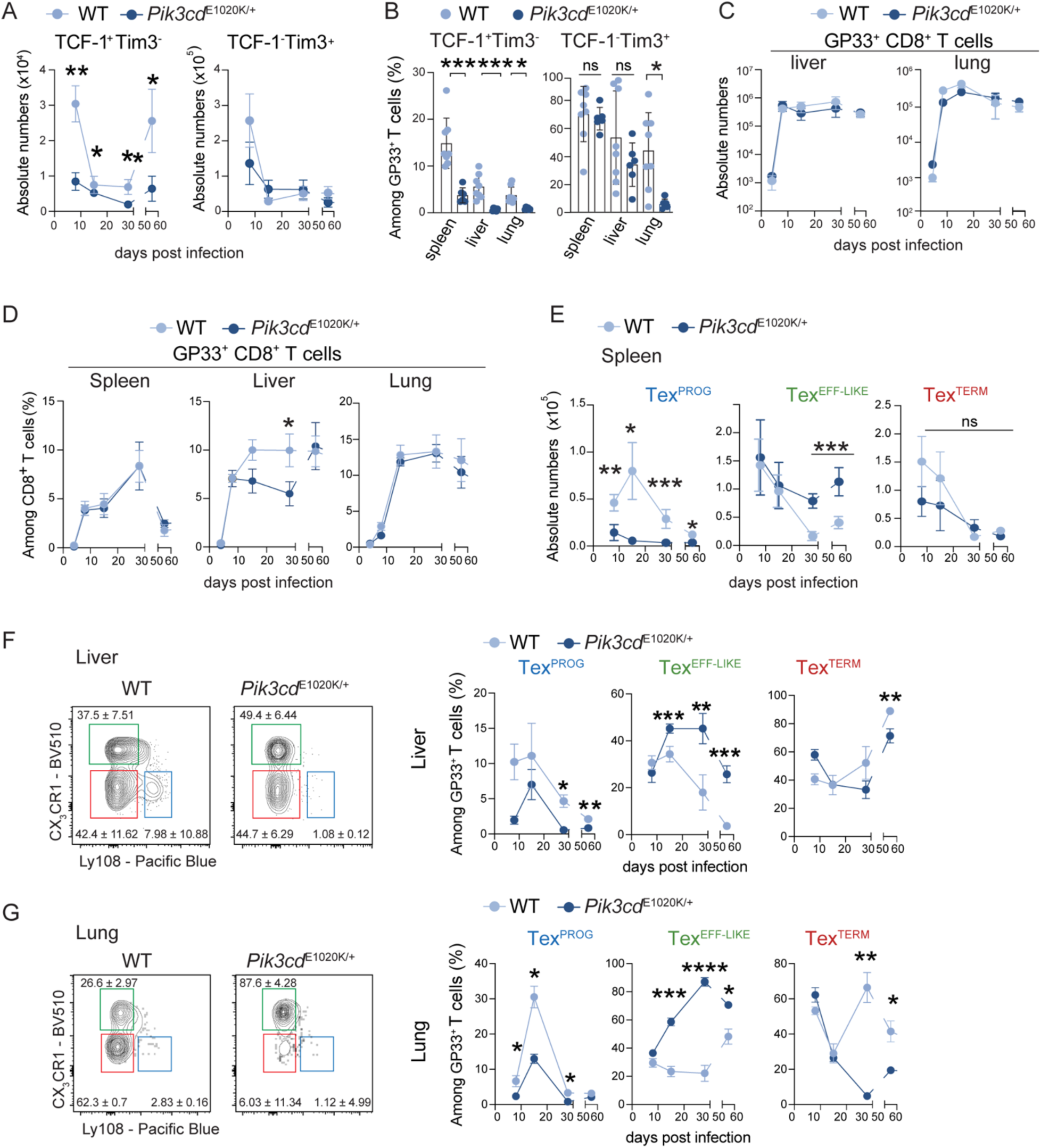
Activated PI3Kδ promotes expansion of CX_3_CR1^+^ effector-like Tex cells. C57BL/6 WT and *Pik3cd*^E1020K/+^ mice were infected i.v. with 2x10^6^ pfu of LCMV cl13. **(A)** Absolute numbers of among GP33^+^ CD8^+^ TCF-1^+^ Tim3^-^ and TCF-1^-^ Tim3^+^ cells on indicated timepoints in spleen. **(B)** Percentages of TCF-1^+^ Tim3^-^ and TCF-1^-^ Tim3^+^ cells among GP33^+^ CD8^+^ T cells at d28 p.i. in indicated organ. **(C)** Absolute numbers of GP33^+^ CD8^+^ T cells in liver (left) and lung (right). **(D)** Percentage of GP33^+^ CD8^+^ T cells on indicated timepoints in spleen (left), liver (middle) and lung (right). **(E)** Absolute number of GP33^+^ CD8^+^ Ly108^+^ CX_3_CR1^-^Tex^PROG^, Ly108^-^ CX_3_CR1^-^ Tex^TERM^ and Ly108^-^ CX_3_CR1^+^ Tex^EFF-LIKE^ cells on indicated timepoints in spleen. **(F-G)** Representative flow plots from d28 p.i. (left) and percentages (right) of Tex^PROG^, Tex^TERM^ and Tex^EFF-LIKE^ cells among GP33^+^ CD8^+^ T cells on indicated timepoints from liver **(F)** and lung **(G).** Data are representative of a pool of at least two independent experiments. Numbers in contour and line plots indicate means ± SEM of shown experiment. In **(A), (C-E)** and **(F-G)** for WT condition, each dot represents a pool of 6, 6, 8 and 11 mice at d8, d15, d28, d55 p.i. respectively, and for *Pik3cd*^E1020K/+^ each dot represents a pool of 8, 8, 6 and 9 mice, at d8, d15, d28, d55 p.i. respectively. In **(B)** each dot represents one mouse. Statistical differences were determined by one-way ANOVA (**A, C-G)** and Student’s t-test **(B)** *p < 0.05, **p < 0.01, ***p < 0.001, ****p <0.0001.

**Extended Data Figure 3.**
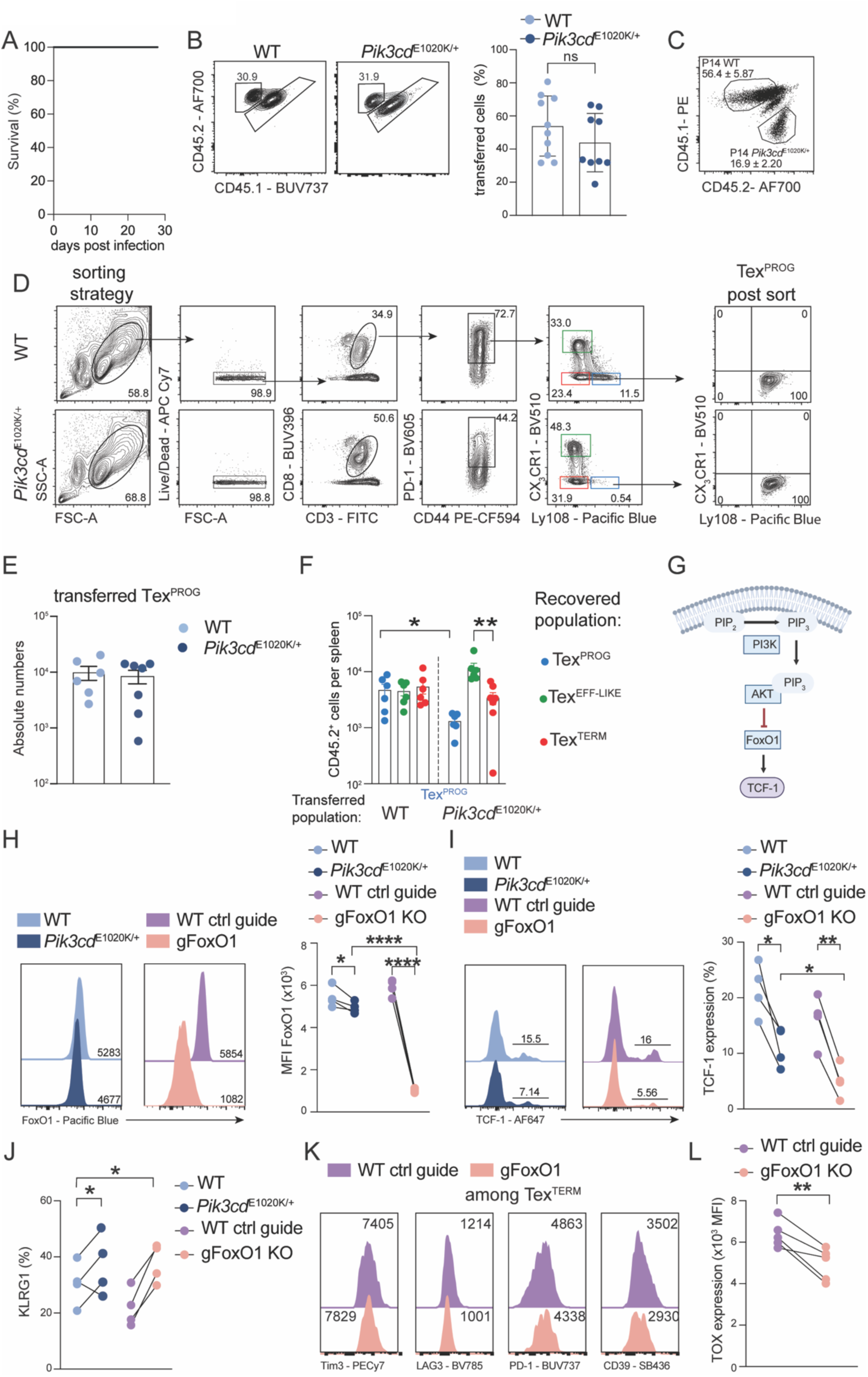
Loss of Tex^PROG^ in activated PI3Kδ mice is FoxO1 dependent. **(A)** CD8^+^ T cells sorted from C57BL/6 P14 (CD45.1) mice and *Pik3cd*^E1020K/+^ P14 (CD45.2) mice were co-transferred into congenic C57BL/6 (CD45.1/2) mice, which were infected with LCMV cl13. Kaplan-Meyer plot showing percentage of survival. **(B)** CD8^+^ T cells sorted from C57BL/6 P14 mice or *Pik3cd*^E1020K/+^ P14 (CD45.2) mice were transferred into congenic C57BL/6 (CD45.1/2) mice, which were infected i.v. with 2x10^6^ PFU of LCMV cl13. Transferred cells were analyzed d8 p.i.. Representative FACS plots (left) and graph (right) showing percentages of transferred cells. **(C)** CD8^+^ T cells sorted from C57BL/6 P14 (CD45.1) mice and *Pik3cd*^E1020K/+^ P14 (CD45.2) mice were co-transferred into congenic C57BL/6 (CD45.1/2) mice, which were infected with LCMV cl13.. Representative FACS plot of transferred populations at d8 p.i.. **(D-E**) Ly108^+^CX_3_CR1^-^ Tex^PROG^ CD8^+^ T cells were sorted from d15 cl13-infected C57BL/6 WT (CD45.1) and *Pik3cd*^E1020K/+^ (CD45.2) mice and transferred into infection-matched congenic C57BL/6 (CD45.1/2) mice. Transferred cells were analyzed d8 p.i.. **(D)** Sorting strategy and post sort analysis of Tex^PROG^ cells. **(E)** Absolute numbers of donor cells d8 post transfer. **(F)** Absolute numbers of transferred Tex^PROG^ cells in each subpopulation. **(G)** PI3K driven activation of AKT leads to phosphorylation, nuclear exclusion and degradation of FoxO1. One target gene of FoxO1 is *Tcf7*. **(H-L)** CD45.1 CRISPR-Cas9 control and CD45.2 CRISPR-Cas9 FoxO1-deficient P14 CD8^+^ T cells were co-transferred into congenic C57BL/6 WT (CD45.1/2) mice, which were infected i.v. with 2x10^6^ PFU of LCMV cl13. Transferred cells were analyzed d8 p.i.. Representative histograms (left) and graphs (right) showing MFI of FoxO1 **(H)** and percentages of TCF-1 **(I)** among transferred cells. **(J)** KLRG1 positive cells among transferred cells. **(K)** Representative histograms showing expression of indicated marker among transferred Tex^TERM^ P14 cells. **(L)** Graph showing TOX MFI of transferred P14 cells. Data are representative of at least two independent experiments, with at least 4 mice/group. Numbers in contour plots indicate percentage. Numbers in histograms indicate MFI **(H and K)** or percentage **(I)**. In **(C)**, **(E-F)** and **(H-L)** one dot represents 1 mouse. Statistical differences were determined by paired two-tailed *t* tests **(H-L)** and Student’s t-test **(C, E)** *p < 0.05, **p < 0.01, ***p < 0.001.

**Extended Data Figure 4.**
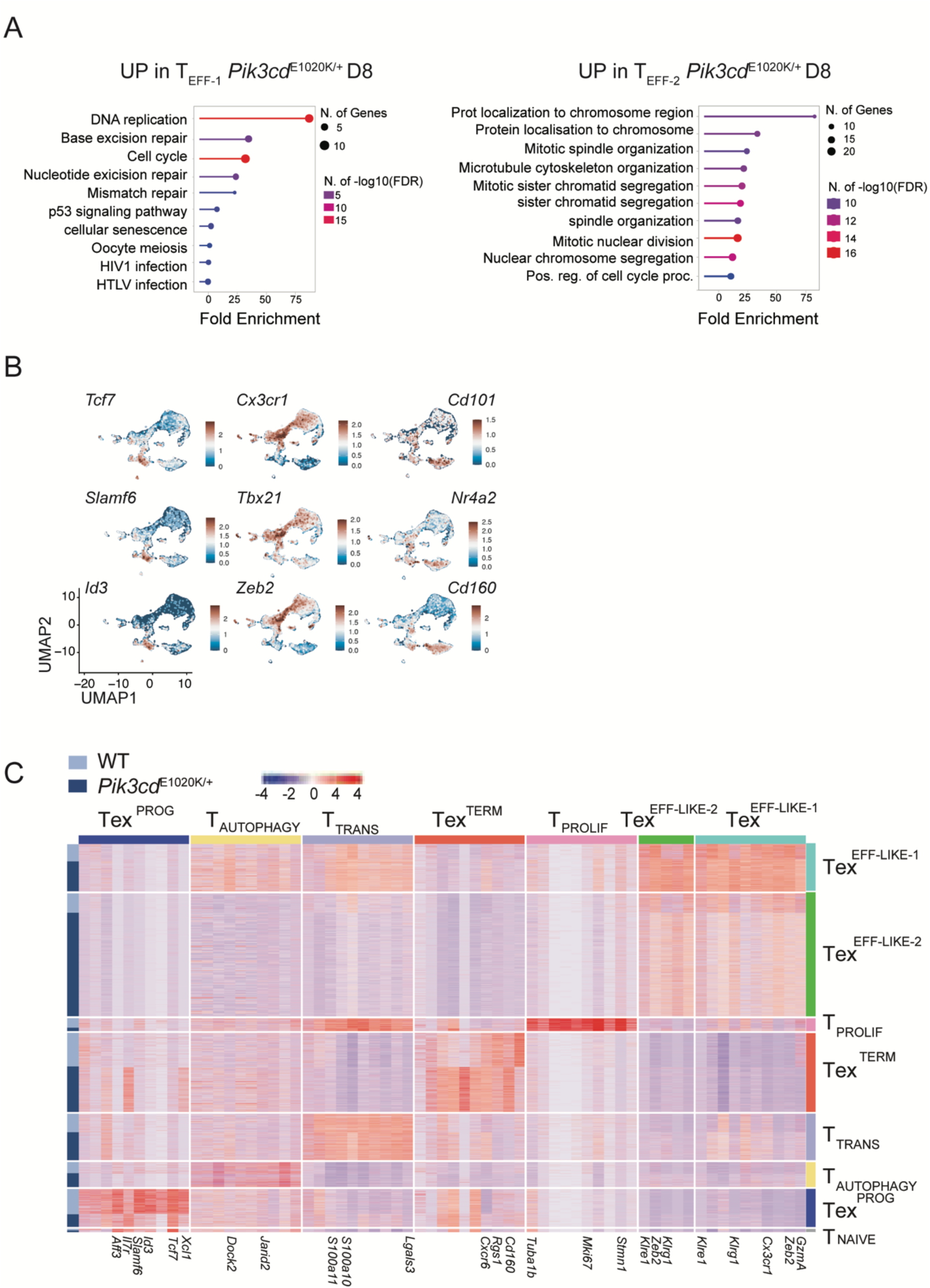
scRNAseq reveals early Tex^EFF-LIKE^ differentiation and expansion in activated PI3Kδ mice. C57BL/6 WT and *Pik3cd*^E1020K/+^ mice were infected i.v. with 2x10^6^ PFU of LCMV cl13 and GP33^+^ CD8^+^ T cells were analyzed and sorted on d8 and d28 p.i. for single cell RNAseq. **(A)** GO pathway analysis on top upregulated DEG in indicated cluster from d8 *Pik3cd*^E1020K/+^ GP33^+^ cells compared to WT. **(B)** Feature plots showing indicated representative genes for progenitor, effector and terminally exhausted cells. **(C)** Heatmap representing the average aggregated expression of top 10 DEGs among the indicated clusters. Color scale indicates *Z* score of gene expression.

**Extended Figure 5.**
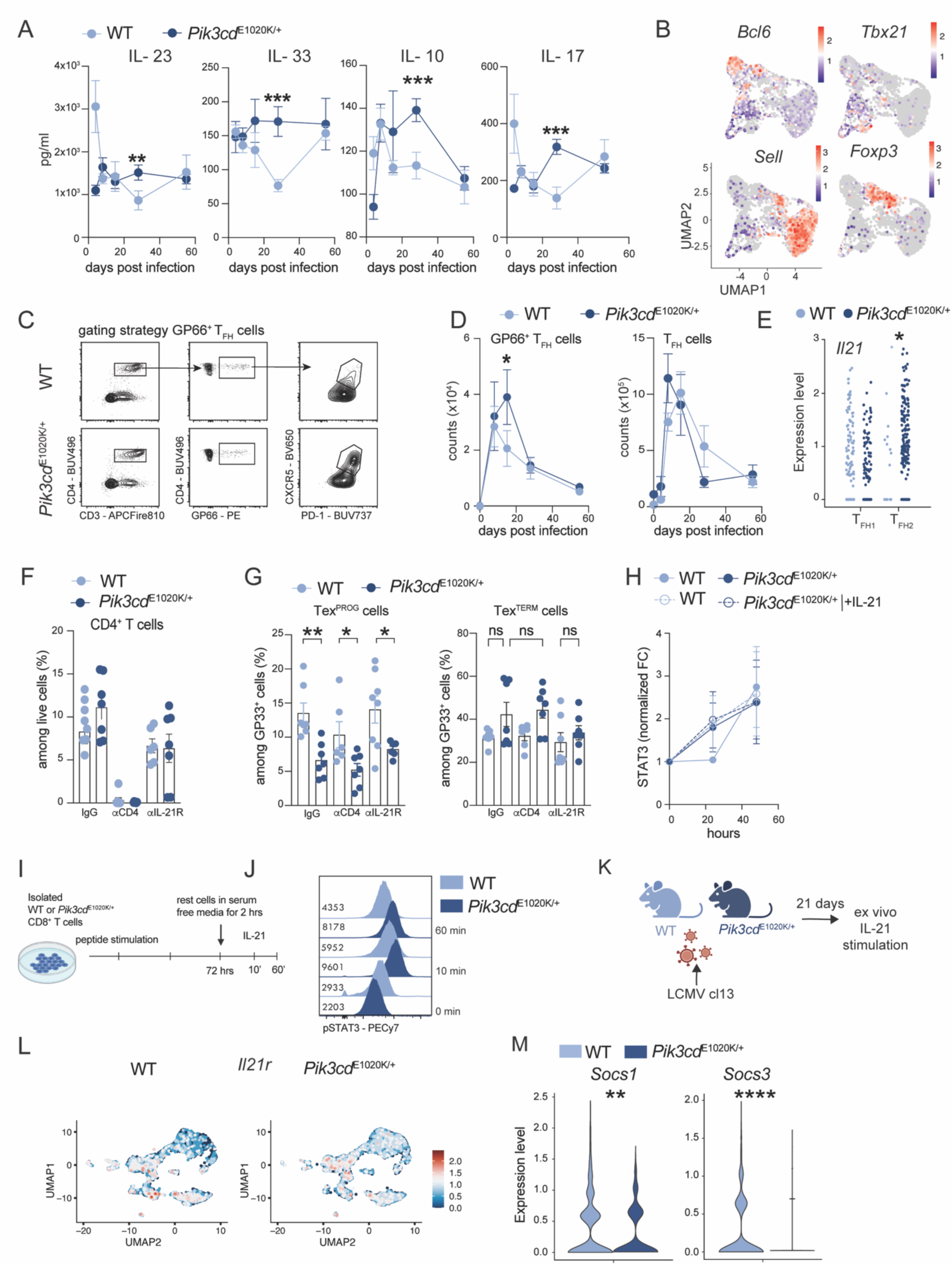
CX_3_CR1^+^ Tex^EFF-LIKE^ cell expansion and persistence is CD4 and IL-21 dependent. **(A-E)** C57BL/6 WT and *Pik3cd*^E1020K/+^ mice were infected i.v. with 2x10^6^ PFU of LCMV cl13. **(A)** Cytokines in liver homogenates at indicated timepoints. **(B)** Features plots of CD4^+^ scRNAseq data of indicated gene. **(C)** Gating strategy of CD4^+^GP66^+^ PD1^+^CXCR5^+^ T_FH_ cells. **(D)** Absolute numbers of indicted population. **(E)** Expression levels of *Il21* in cells among indicated cluster. (**F-G)** C57BL/6 WT and *Pik3cd*^E1020K/+^ mice were infected with LCMV cl13. Mice were either treated with αCD4 or αIL-21R mAbs and analyzed d8 p.i.. **(F)** Percentages of CD4^+^ T cells. **(G)** Percentages of Tex^PROG^ (left) and Tex^TERM^ (right) cells among GP33^+^ T cells. **(H)** CD8^+^ T cells were stimulated *in vitro* with anti-CD3/28 mAbs and IL-21 for 3 days and analyzed. Fold change of total STAT3. **(I-J)** OT-1 CD8^+^ T cells were activated with OVA_257-264_ peptide for 3 days, then rested 2 hrs in serum free media and re-stimulated with IL-21 for 10 and 60 mins. **(I)** Experimental design and **(J)** representative histogram of pSTAT3 MFI. **(K)** CD8^+^ T cells isolated from d21 LCMV cl13 infected C57BL/6 WT and *Pik3cd*^E1020K/+^ mice were stimulated *ex vivo* for 2 hrs with IL-21 and analyzed by flow cytometry. Experimental design for Fig 5P. **(L)** Feature plot from CD8 d28 p.i. scRNAseq data **(see Fig.4E)** showing *Il21r* levels. **(M)** Violin plots showing expression levels of *Socs1* and *Socs3* from d28 p.i. scRNAseq data. In **(A, D, F, G, H)**, data shows pool of at least two independent experiments. In **(J)**, data is representative of three independent experiments In **(A)** in WT condition, each dot represents a pool of 4, 8, 4, 8, 4 mice at d4, d8, d15, d28 and d55 p.i. respectively and in *Pik3cd*^E1020K/+^ condition, each dot represents a pool of 4, 7, 4, 8, 4 mice at d4, d8, d15, d28 and d55. In **(D)** in WT condition, each dot represents a pool of 6, 6, 7, 7, 9 mice at d4, d8, d15, d28 and d55 p.i. respectively and in *Pik3cd*^E1020K/+^ condition, each dot represents a pool of 6, 7, 6, 7, 9 mice at d4, d8, d15, d28 and d55. In **(E)** one dot represents one cell. In **(F-G),** one dot represents one mouse. In **(H)** each dot represents the mean of one experiment. Statistical differences were determined by one-way ANOVA (A, D, F-G, H) *p < 0.05, **p < 0.01, ***p < 0.001, ****p <0.0001.

**Extended Data Figure 6.**
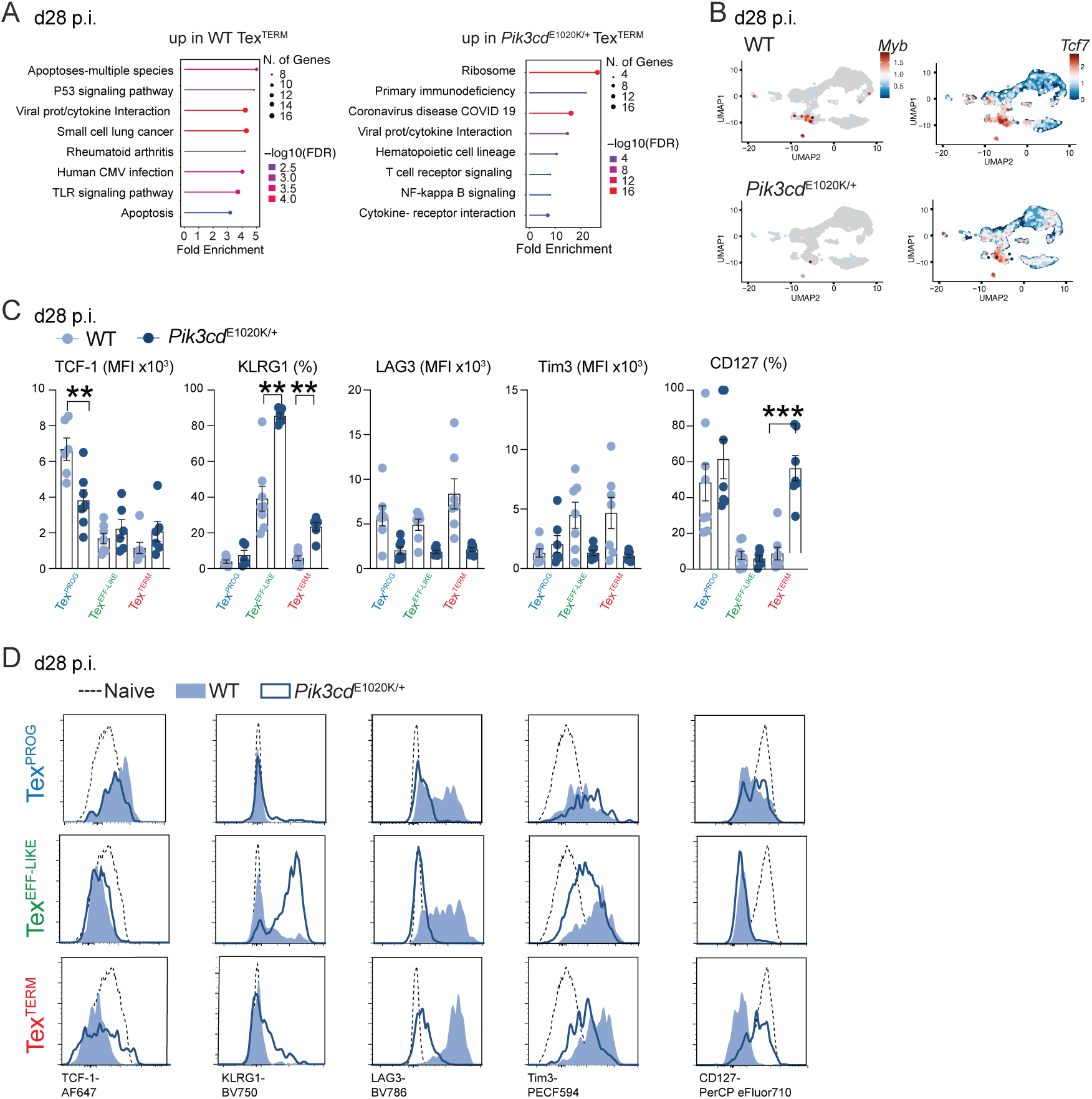
Activated PI3Kδ suppresses TOX and exhaustion program. **A)** GO pathway analysis on top upregulated DEGs (p > 0.05) in WT and *Pik3cd*^E1020K/+^ Tex^TERM^ cells from d28 p.i. scRNAseq data**. (B)** Feature plots showing *Myb* (left) and *Tcf7* (right) expression at d28 p.i.. **(C-D)** Graphs **(C)** and representative histogram **(D)** showing percentages or MFI of indicated marker. In **(C)** each dot represents one mouse, shown is a pool of two independent experiment, with 3-4/group. Statistical differences were determined Student’s t-test **(C)** *p < 0.05, **p < 0.01, ***p < 0.001, ****p <0.0001.

**Extended Data Figure 7.**
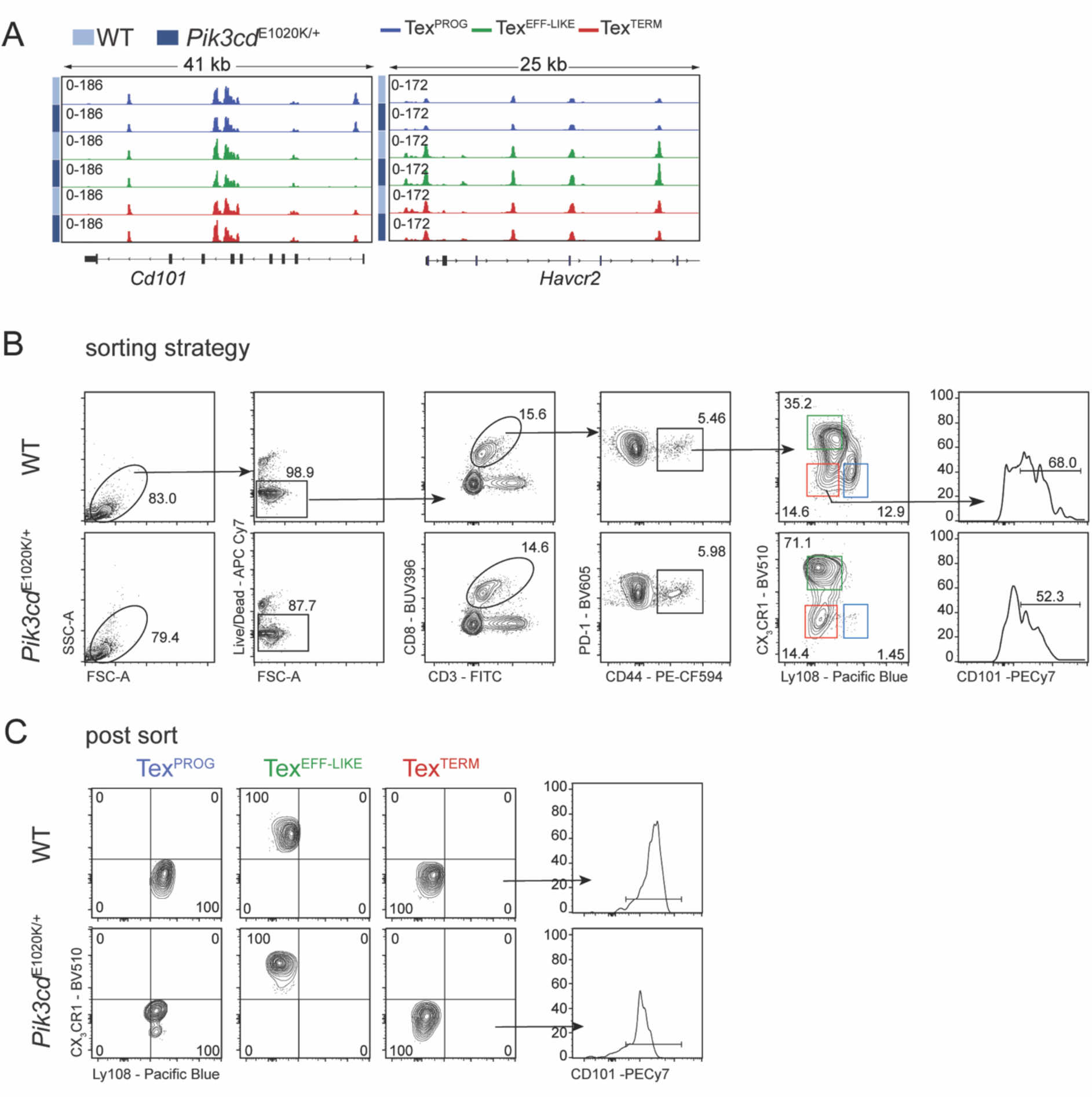
Activated PI3Kδ prevents changes associated with exhaustion and promotes plasticity. **(A)** GP33^+^ antigen-specific Tex^PROG^, Tex^EFF-LIKE^ and Tex^TERM^ cells were sorted from C57BL/6 WT or *Pik3cd*^E1020K/+^ mice d15 p.i. with LCMV cl13 and analyzed by ATAC-seq. ATAC-seq tracks at selected genes (*Cd101 and Havcr2*) from Tex^PROG^, Tex^EFF-LIKE^ and Tex^TERM^ cells. **(B-C)** D15 p.i. sorted Tex^TERM^ cells from C57BL/6 WT and *Pik3cd*^E1020K/+^ mice were transferred into infection matched host. Representative FACS plots showing sorting strategy **(B)** and post sort analysis **(C)**. Numbers in contour plot indicate percentage.

